# Optimizing alkalinity control in Recirculating Aquaculture Systems (RAS): a dynamic modelling approach

**DOI:** 10.1101/2024.06.28.600787

**Authors:** Marie Aline Montjouridès, Susanna Röblitz, Håkon Dahle

**Affiliations:** Computational Biology Unit (CBU), Department of Biology, Thormøhlensgate 55, Bergen, 5008, Norway; Computational Biology Unit (CBU), Department of Informatics, Thormøhlensgate 55, Bergen, 5008, Norway

**Author notes:** corresponding author *Email address:* (Marie Aline Montjouridès).

**Keywords:** dynamic modelling, alkalinity, optimization, RAS

## Abstract

Maintaining good water quality is essential for successful fish production in land-based recirculating aquaculture systems (RAS). Numerous interdependent factors influence water quality parameters, making it difficult to evaluate which operational strategies are most favorable. Mathematical models and model simulations have proven to be powerful tools to evaluate how RAS design and operation are linked to RAS dynamics, but these models rarely implement pH and carbonate species as dynamic variables. Here, we present a dynamic model for RAS (dynRAS) that combines rates of TAN (total ammonia nitrogen) removal, fish growth, and *CO*_2_ and TAN excretion, with a reaction model for pH and the carbonate system formulated based on the law of mass action. A novel aspect of our approach is the incorporation of a dosage system modelled by a Hill-function, enabling the exploration of diverse dosing strategies for pH and alkalinity management. The model was validated based on empirical data from a pilot scale RAS system operated with a feeding regime involving 12 hours of feeding per day. We found that model simulations could be used to accurately predict diurnal cycling patterns in RAS water quality parameters. Furthermore, we made use of simulations to assess how diurnal cycling varies with changing pH and alkalinity levels. Model results emphasize the complexity of pH and alkalinity control in RAS in relation to overall water quality management. Based on our simulations, we argue that what should be considered as optimal pH and alkalinity in RAS depends on the state of the system. Accordingly, optimal pH and alkalinity thresholds may vary between different RAS units and between different time points of a rearing cycle. More generally, we demonstrate how modelling and model simulations can be an effective way of getting insights into the dynamics of complex RAS interactions and provide a valuable tool to efficiently explore effects of different operational strategies on water quality parameters.

## 1. Introduction

Recirculating aquaculture systems (RAS) have the potential to reduce the ecological impact of fish farming by reducing water usage and minimizing waste discharge in the environment [1, 2]. These systems allow inland fish production and reach average water recirculation rates of more than 95% per day [3]. One of the primary advantages of RAS is the ability to grow fish in a controlled environment. However, maintaining suitable water quality parameters in a closed-loop system is challenging given the occurrence of dynamic biological processes, and remains a primary issue in the operation of RAS [4]. Because waste production dynamics are closely tied to fish growth and feeding, these dynamics can change over timescales ranging from minutes to months [5]. These long-term dynamics make it costly and time-consuming to solely rely on experimentation for further water quality management development.

Model simulations provide a complementary approach to experimental trials, particularly for optimizing the control of parameters such as pH and alkalinity that are involved in a multitude of chemical equilibria and biological processes [6, 7, 8]. In RAS, fluctuations in pH can have deleterious effects on the toxicity of metabolites such as *NH*_3_ and *CO*_2_. To reduce pH fluctuations, RAS operators commonly add a base or carbonate buffer in the system to maintain the buffering capacity of the water. This buffering capacity is referred to as alkalinity. More specifically, total alkalinity (TA) refers to the amount of base that can neutralize hydrogen ions (*H*^+^). It can be approximated by eq. (1) because in aquatic systems, alkalinity primarily originates from the carbonate buffer system (eqs. (2),(3)) which involves carbonic acid (*H*_2_*CO*_3_) and its conjugate bases (bicarbonate 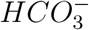 and carbonate 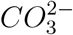 [9]:

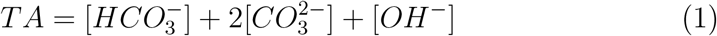

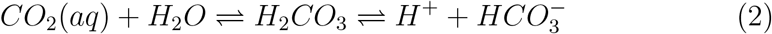

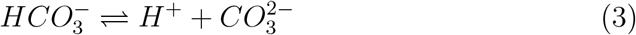

Control strategies for alkalinity are based on this equilibrium, either by raising pH through strong base addition such as *NaOH* or increasing bicarbonate via carbonate buffer addition. If not designed properly, chemical supplementation of *NaOH* or *NaHCO*_3_ can result in excessively high pH or *CO*_2_ levels. While there are common practices for alkalinity control, suggested alkalinity levels range from 50 to 300 mg/L as *CaCO*_3_ depending on the species and the water system used [10, 11]. Considering the complexity of the system dynamics, designing and refining alkalinity control in RAS can be challenging for operators due to the lack of consensus on best practices. Achieving the correct balance between pH, alkalinity, and *CO*_2_ levels is difficult without appropriate decision-support tools.

A dynamic model incorporating processes specific to RAS has already been developed in [5]. This model is based on activated sludge models [12, 13] and integrates fish growth and fish metabolite production. Later refined in [14], the model has proven to be a useful tool for investigating different RAS topologies for optimizing water treatment systems [15]. However, while the model implements a control loop for alkalinity, pH and pH control are not included as dynamic processes. To our knowledge, only one recent publication [16] includes pH in a model for RAS. Using a dynamic model and reaction invariant approach, the authors included pH and alkalinity in two models and tested different configurations for pH and alkalinity adjustment [16]. Yet, both models assume a constant feeding rate and a *CO*_2_ production rate independent of fish size as the modelled dynamics mainly focus on short timescales. Since oxygen consumption and thus *CO*_2_ excretion is dependent on fish size [17], practices for water quality management may need to change throughout a production cycle to ensure optimal rearing conditions. As such, including changes in *CO*_2_ excretion rate within a modelled carbonate system for longer simulations could improve model predictions, providing better insights into how water quality management practices should be adapted over time, especially with regard to alkalinity control.

By implementing a mechanistic model combining the slow biological processes and the rapid chemical reactions of the carbonate system, our work seeks to demonstrate how such a model can be used to evaluate water quality management practices with a focus on alkalinity. This model allows the simulation of different alkalinity levels as well as a pH and alkalinity control system. Total ammonia nitrogen (TAN) and *CO*_2_ excretion production rates related to fish metabolism have been included for a more comprehensive representation of the system’s dynamic. The ammonium dissociation reaction has been included allowing the implementation of *NH*_3_ removal [18] by the ammonia-oxidizing bacteria (AOB), which, to our knowledge, has not been implemented as such in previous RAS models. We use high-resolution measurement data for *CO*_2_, pH and alkalinity from a pilot scale RAS to evaluate the model predictions. Overall, we provide what could be the basis for the development of a simulation tool that operators can use to identify the most effective buffering strategies in coherence with the system’s overall dynamic.

## 2. Method

### 2.1. System configuration

The model was developed based on an experimental set-up (Fig. 1) carried out in pilot-scale RAS units (NanoRas, Alpha Aqua, Denmark) at Marineholmen RASLab (Bergen, Norway) from August 2022 to January 2023 [19]. This unit comprises a 1000 L tank, a swirling operator for solid removal followed by a drum filter, three moving bed biofilter reactor chambers of 320 L each, and a degasser compartment of about 700 L. Two peristatic pumps were displayed after the tank upstream of the biofilter and were dosing either *NaOH* or *NaHCO*_3_.

**Fig. 1.**
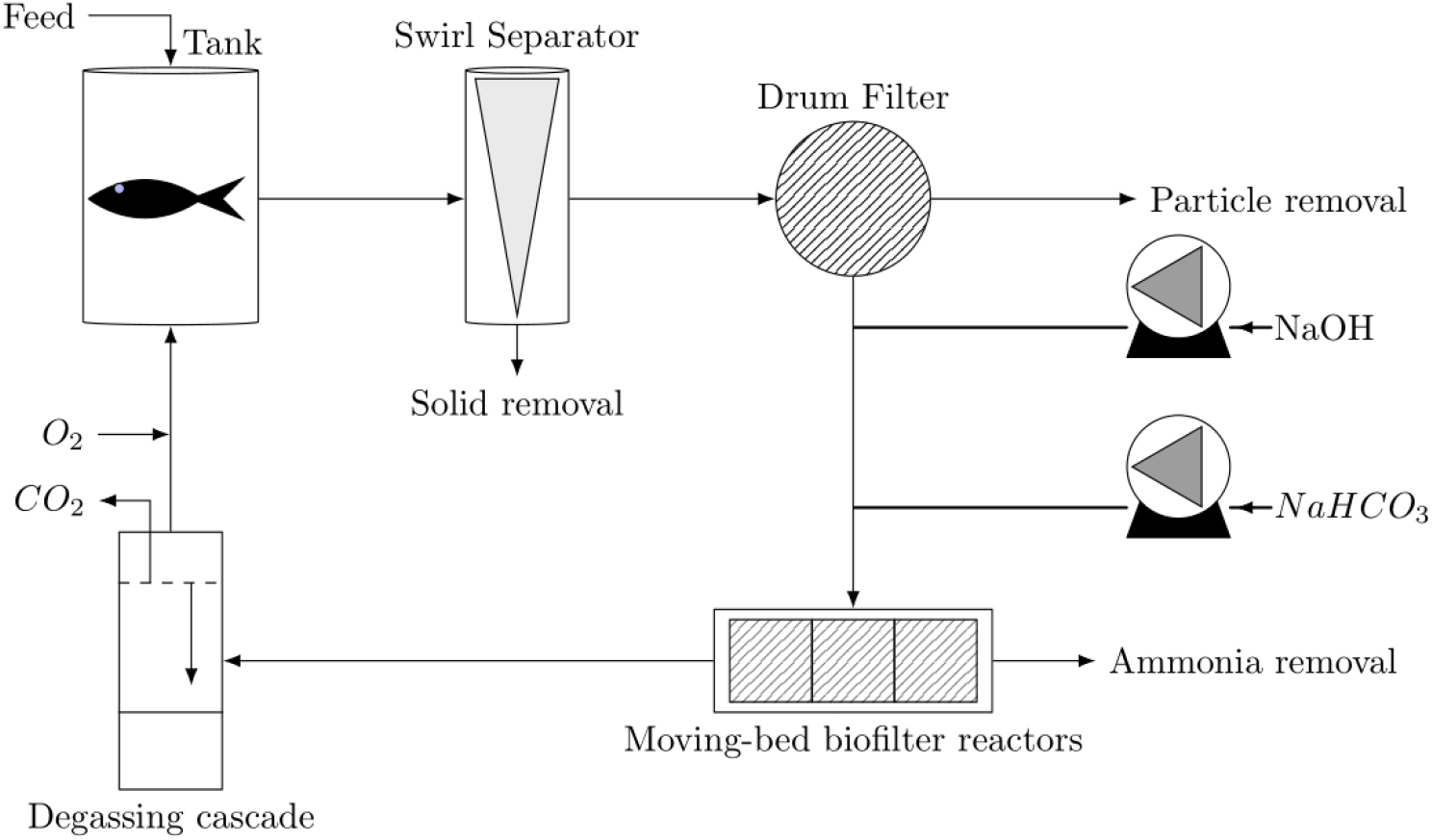
Principal compartments of a RAS. Two peristaltic pumps were placed upstream of the biofilter in the context of the experimental trial to control akalinity and pH with *NaHCO*_3_ and *NaOH*.

### 2.2. Data collection

A continuous monitoring system (AquaSENSE™, SEARAS AS, Bergen, Norway [20]) was placed between the tank and the bio-filter to record *CO*_2_, pH, and alkalinity levels of the water coming from the tank. In addition, as no continuous monitoring systems for TAN were available at the time of the experimental trial, samples for TAN measurement were collected at the same location for continuous monitoring. The measurements were taken every two weeks with samples taken every 2 hours (h) throughout the feeding period. One additional sampling point was taken on the following day 1 h prior to the start of the next feeding period. The TAN samples were analyzed through spectrophotometry using a Spectroquant® Pharo 300 spectrophotometer (Spectroquant® Pharo 300 spectrophotometer, MERCK, Darmstadt, Germany) and ammonium-Test (Supelco Spectroquant® Ammonium-Test, MERCK, Darmstadt,Germany).

### 2.3. Model implementation

The dynRAS model is a dynamic model implemented in Python 3.8.18 using the Scipy library for the solution of systems of ordinary differential equations (ODE). dynRAS was developed using an object-oriented programming approach (Fig. 2) making the model flexible to modifications and changes in system topology. The principal components implemented in the model are the fish tank, the biofilter, and the degasser. The removal of solids and particles is not included in the model and the biofilter chambers are modelled as one large stirred compartment with the assumption of a homogeneous medium. For simplicity, oxygen is considered saturated. Although the parameters used were determined for a temperature of 14 °C and salinity of 35 ppt, the relative dependencies for the system species would be consistent across different temperatures and salinity. However, for improved quantitative accuracy, the equilibrium constants and the thermal growth coefficient (TGC) should be adjusted accordingly. Each of the system compartments includes the same set of chemical species in *mmol* · *L*^−1^(see Tab. 1) with their specific biological or mechanical processes (Fig. 2). Transport of chemical species between components occurs via a defined flow rate in *L* · *s*^−1^, denoted as *F*. Water exchanges are modelled in the degasser with incoming flows from a source compartment and flow out of the system. The exchange flow (*F*_*ex*_) is calculated based on the total volume of the system and the percentage of water recirculating per day. Fixed concentrations for all the species from the external source are set to 0 except for *H*^+^, *OH*^−^, 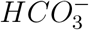, and 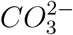 where we used fixed concentrations of 10^−4.5^, 10^−3.5^, 1.15 and 10^−3^ *mmol* · *L*^−1^, respectively, to simulate a small external addition of 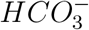 to the system (Fig. 2).

**Fig. 2.**
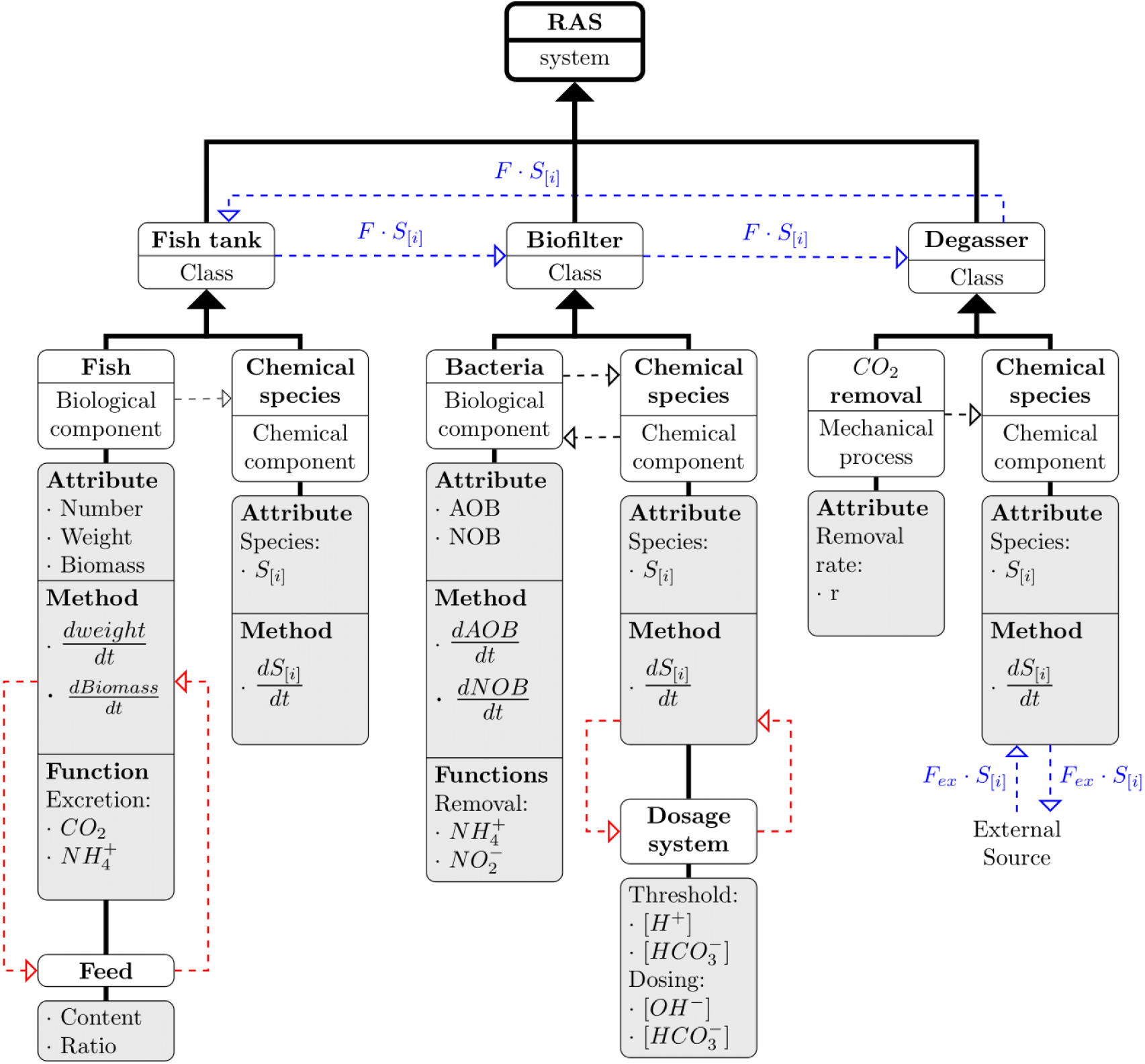
Schematic of the modelling workflow. Red dashed arrows represent feedback-loops implemented in the model, blue dashed arrows represent the flow of chemical species between the classes, and black dashed arrows represent interactions between components. Each of the represented classes possesses attributes, methods, and functions relative to its specific compartment. The full expression of the ODEs in each compartment can be found in Appendix B.

**Table 1:** Overview of sytem variables in the Fish Tank (T), Biofilter (B), and the Degasser (D). The check mark and cross mark denote the presence or absence of the variable in the compartment. *S*_*i*_ denotes the chemical species *i* in *mmol* · *L*^−1^.

| Variable | Description | Unit | T | B | D |
| --- | --- | --- | --- | --- | --- |
| $S_{\text{CO}_{2\text{aq}}}$ | Carbon dioxide concentration | $\text{mmol} \cdot \text{L}^{-1}$ | ✓ | ✓ | ✓ |
| $S_{\text{HCO}_3^-}$ | Bicarbonate concentration | $\text{mmol} \cdot \text{L}^{-1}$ | ✓ | ✓ | ✓ |
| $S_{\text{CO}_3^{2-}}$ | Carbonate concentration | $\text{mmol} \cdot \text{L}^{-1}$ | ✓ | ✓ | ✓ |
| $S_{\text{H}^+}$ | Protons concentration | $\text{mmol} \cdot \text{L}^{-1}$ | ✓ | ✓ | ✓ |
| $S_{\text{OH}^-}$ | Hydroxide ions concentration | $\text{mmol} \cdot \text{L}^{-1}$ | ✓ | ✓ | ✓ |
| $S_{\text{NH}_4^+}$ | Ammonium concentration | $\text{mmol} \cdot \text{L}^{-1}$ | ✓ | ✓ | ✓ |
| $S_{\text{NH}_3}$ | Ammonia concentration | $\text{mmol} \cdot \text{L}^{-1}$ | ✓ | ✓ | ✓ |
| $S_{\text{NO}_2^-}$ | Nitrite concentration | $\text{mmol} \cdot \text{L}^{-1}$ | ✓ | ✓ | ✓ |
| AOB | Biomass ammonia-oxidizing bacteria | mg | ✗ | ✓ | ✗ |
| NOB | Biomass nitrite-oxidizing bacteria | mg | ✗ | ✓ | ✗ |
| Weight | Fish weight | g | ✓ | ✗ | ✗ |
| Biomass | Fish Biomass | g | ✓ | ✗ | ✗ |

### 2.4. Model construction and process description

The dynRAS model integrates the main processes occurring in a RAS: fish growth, TAN and *CO*_2_ excretion and removal. These processes are combined with a reaction model describing the chemical reactions associated with the carbonate system. The following sections describe the model construction and processes included in the model.

#### 2.4.1. Reaction model

To simulate the variations in water chemistry within the system, a reaction model was formulated based on the mass action rate law. This model is grounded in the kinetics of the carbonate system, following an approach similar to methods described in [21]. Reactions (2) and (3) can be reformulated as reactions (4) and (5), where *k* and *k*_−_ correspond to the forward and backward rate constants of the reaction, respectively. To adequately model pH variations, reaction (6) for the ionization of water needs to be included. In addition, we included the ammonium-to-ammonia dissociation reaction (7) to model the TAN distribution within the system, distinguishing between the ammonia and ammonium forms.

##### CO_2_ to 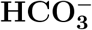

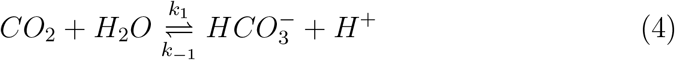

While the reaction between *H*_2_*CO*_3_ and 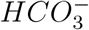 can be considered instantaneous, the reaction of hydration for *CO*_2_ is relatively slow (seconds) [21, 22]. This kinetic results in most of the *CO*_2_ present in the system occurring in the form of *CO*_2_ and the fraction of *H*_2_*CO*_3_ in the system is very small. Therefore, reaction (4) is a simplification of reaction (2) where the notation *CO*_2_ denotes the sum of *CO*_2_ aqueous and *H*_2_*CO*_3_ in the system.

##### 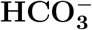 dissociation to 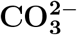

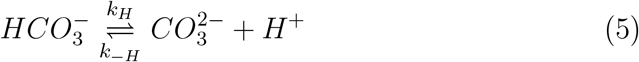

##### Water Ionization

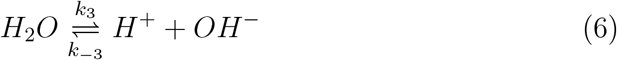

##### Ammonium to Ammonia dissociation reaction

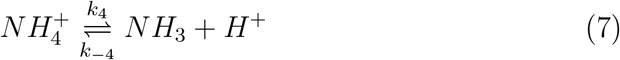

The system of differential equations was formulated according to the mass action rate law. To illustrate the application of this law, we can consider the following reaction,

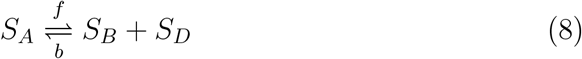

where *S*_*A*_ denotes the reactant, *S*_*B*_ + *S*_*D*_ is the product of the reaction, and *f* and *b* are the forward and backward rate constant. The rate of consumption *r*_*C*_ and production *r*_*P*_ of *S*_*A*_ can be formulated as:

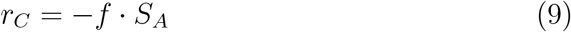

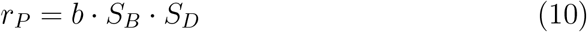

Note that the species names in these two equations and onward also represent the species concentrations. The species concentrations are time-dependent but for improved readability, time is omitted in the following. The overall variation of *S*_*A*_ over time can thus be generalized given by the sum of the production and consumption rates:

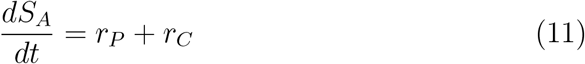

Following this approach, the overall dynamic of each species *S*_*i*_ in our system can be described by the following general formula

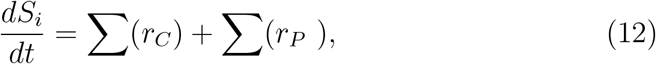

where *i* denotes the species (Tab. 1). Applying this to each species in the system and based on reactions (4-7), species dynamics are as follows:

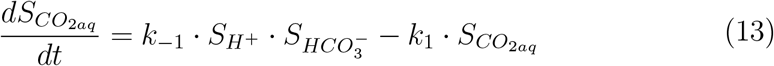

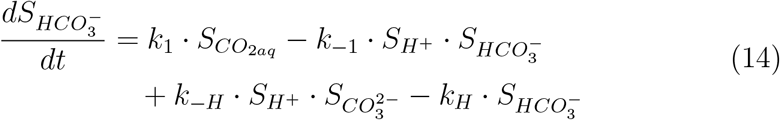

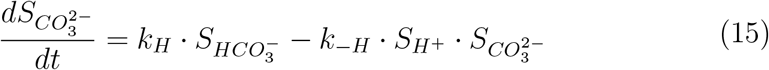

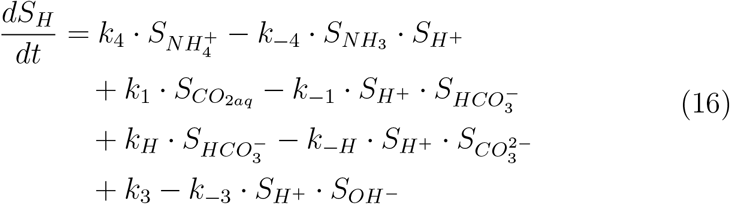

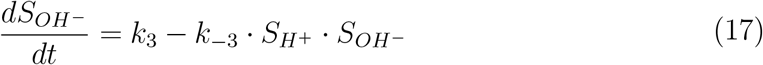

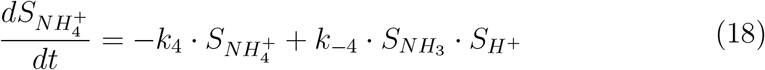

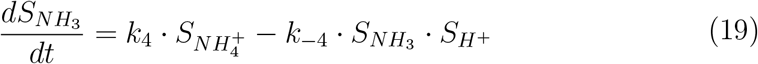

Parameters for equations (12) to (18) can be found in Tab. 2.

**Table 2:** List of the parameters used for the simulation. Most of the parameters for the carbonate system were extracted from the literature [21, 24, 31]. Backward rate constants *k*_−*H*_, *k*_−3_, and *k*_−4_ were recalculated using equilibrium constants at 15°C [32, 30] as we could not find data for 14°C. The carrying capacity of the bio-filter was approximated by weighing biomass extracted from bio-carriers sampled during the experimental trial of Jafari et al.,(2024) [19] and assuming that 5% of that biomass can be allocated to NOB and AOB in the biofilter.

| Parameter | Value | Unit | Description | Reference |
| --- | --- | --- | --- | --- |
| F | 1.0 | L.s<sup>-1</sup> | Flow rate between compartments | - |
| $k_1$ | $1.49 \times 10^{-2}$ | s<sup>-1</sup> | Forward rate constant for $CO_2$ dissociation to $HCO_3^-$ | [24] |
| $k_{-1}$ | 18.9 | L.mmol<sup>-1</sup>.s<sup>-1</sup> | Backward rate constant for $CO_2$ dissociation to $HCO_3^-$ | [24] |
| $k_H$ | 25.059 | s<sup>-1</sup> | Forward rate constant for $HCO_3^-$ dissociation to $CO_3^{2-}$ | [21], recalculated for 15°C |
| $k_{-H}$ | $5.0 \times 10^7$ | L.mmol<sup>-1</sup>.s<sup>-1</sup> | Backward rate constant for $HCO_3^-$ dissociation to $CO_3^{2-}$ | [21] |
| $k_3$ | 1.4 | mmol.L<sup>-1</sup>.s<sup>-1</sup> | Forward rate constant for water ionisation | [21] |
| $k_{-3}$ | $3.11 \times 10^8$ | L<sup>-1</sup>.mmol<sup>-1</sup>.s<sup>-1</sup> | Backward rate constant for water ionisation | [21], recalculated for 15°C |
| $k_4$ | 11.739 | s<sup>-1</sup> | Forward rate constant for $NH_4^+$ dissociation to $NH_3$ | [31], calculated for 15°C |
| $k_{-4}$ | $4.3 \times 10^7$ | L.mmol<sup>-1</sup>.s<sup>-1</sup> | Backward rate constant for $NH_4^+$ dissociation to $NH_3$ | [31] |
| $\mu_{AOB}$ | $6.08 \times 10^{-6}$ | s<sup>-1</sup> | Maximum growth rate for AOB | [14] |
| $\mu_{NOB}$ | $7.64 \times 10^{-6}$ | s<sup>-1</sup> | Maximum growth rate for NOB | [14] |
| $K_{NH3}$ | $7.14 \times 10^{-4}$ | mmol.L<sup>-1</sup> | Half saturation constant for $NH_3$ | Estimated from $K_{NH}$ from [14] |
| $K_{Alk}$ | 0.3 | mmol.L<sup>-1</sup> | Half saturation constant for $HCO_3^-$ | [14] |
| $K_{NO2^-}$ | 0.0714 | mmol.L<sup>-1</sup> | Half saturation constant for $NO_2^-$ | [14] |
| $\rho_{AOB}$ | $1.16 \times 10^{-6}$ | s<sup>-1</sup> | Decay rate for AOB | [14] |
| $\rho_{NOB}$ | $1.16 \times 10^{-6}$ | s<sup>-1</sup> | Decay rate for NOB | [14] |
| $Y_{AOB}$ | 2.94 | g-biomass/moles-NH3 | Biomass Yield | [14] |
| $Y_{NOB}$ | 0.56 | g-biomass/moles-$NO_2^-$ | Biomass Yield | [14] |
| B | 1.2 | kg | Total carrying capacity of the bio-filter for AOB and NOB | Adapted for the model, approximated from biomass and DNA extraction (data not shown) |
| T | 14 | °C | Temperature | - |
| w0 | 70 | g | Initial Fish weight | - |
| TGC | 2.7 | - | Thermal growth rate coefficient | [25, 14] |
| FR | 2.7 | % | Feed Ratio 2.7 % of the body weight | - |
| PC | 46 | % | Protein content of the Feed | Skretting Nutra RC 3 |
| $\alpha$ | 3 | - | $\alpha$ parameter of the hill function | Adapted for the model |
| $\beta$ | 3 | - | $\beta$ parameter of the hill function | Adapted for the model |
| n | 10 | - | Hill coefficient for the dosage system | Adapted for the model |
| r | 0.0015 | s<sup>-1</sup> | $CO_2$ removal rate constant | Adapted for the model |

#### 2.4.2. Dosage Control

In addition to the chemical dynamic, a dosage system is implemented within the biofilter compartment. The dosage system is designed to control *H*^+^ and/or 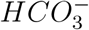 concentration using either sodium hydroxide (*NaOH*), modelled by the addition of *OH*^−^, or using bicarbonate 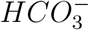 modelled by the addition of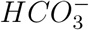. The framework of the dosage system is implemented on a structure akin to a Hill function [23] integrated into a feedback loop enabling dynamic adaptability to fluctuating conditions and pH levels. The dosage functions for the addition of *OH*^−^ and 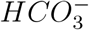 at a time *t* are defined as follows,

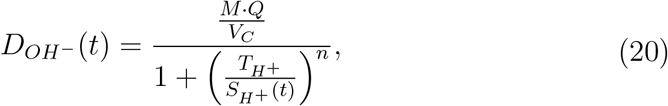

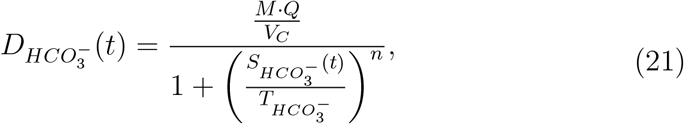

where 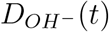 is the dosage system for the addition of *OH*^−^ and 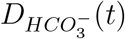 the dosage system for the addition of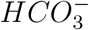. *M* represents the concentration of the dosing agent, *Q* is the flow rate from the dosing compartment to the targeted compartment, *V*_*C*_ is the volume of the compartment receiving the dosage and *t* is the time in seconds.

For pH control, the system is designed to respond to the concentration of *H*^+^ 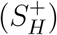 in relation to a threshold 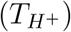. As the *H*^+^ concentration approaches or exceeds 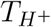, the dosage of the base increases (Fig. 3a). In contrast, when the *H*^+^ concentration falls below the threshold, the dosage is gradually reduced (Fig. 3a). In the case of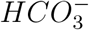, the operational strategy is converse (Fig. 3a). A 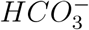 concentration 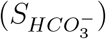 below its respective threshold 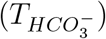 leads to an increase in the concentration of the dosing chemical within the system (Fig. 3b). This increment continues until the 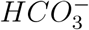 concentration reaches the threshold, after which a decrease in the dosing chemical ensues if the threshold is exceeded (Fig. 3b). The maximum dosing rate is thus determined by the fraction 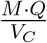.While the dosing rate can be very small it will never reach zero. This allows continuity of the dosage system, avoiding numerical instabilities that were encountered when the dosage was implemented as a discrete event.

**Fig. 3.**
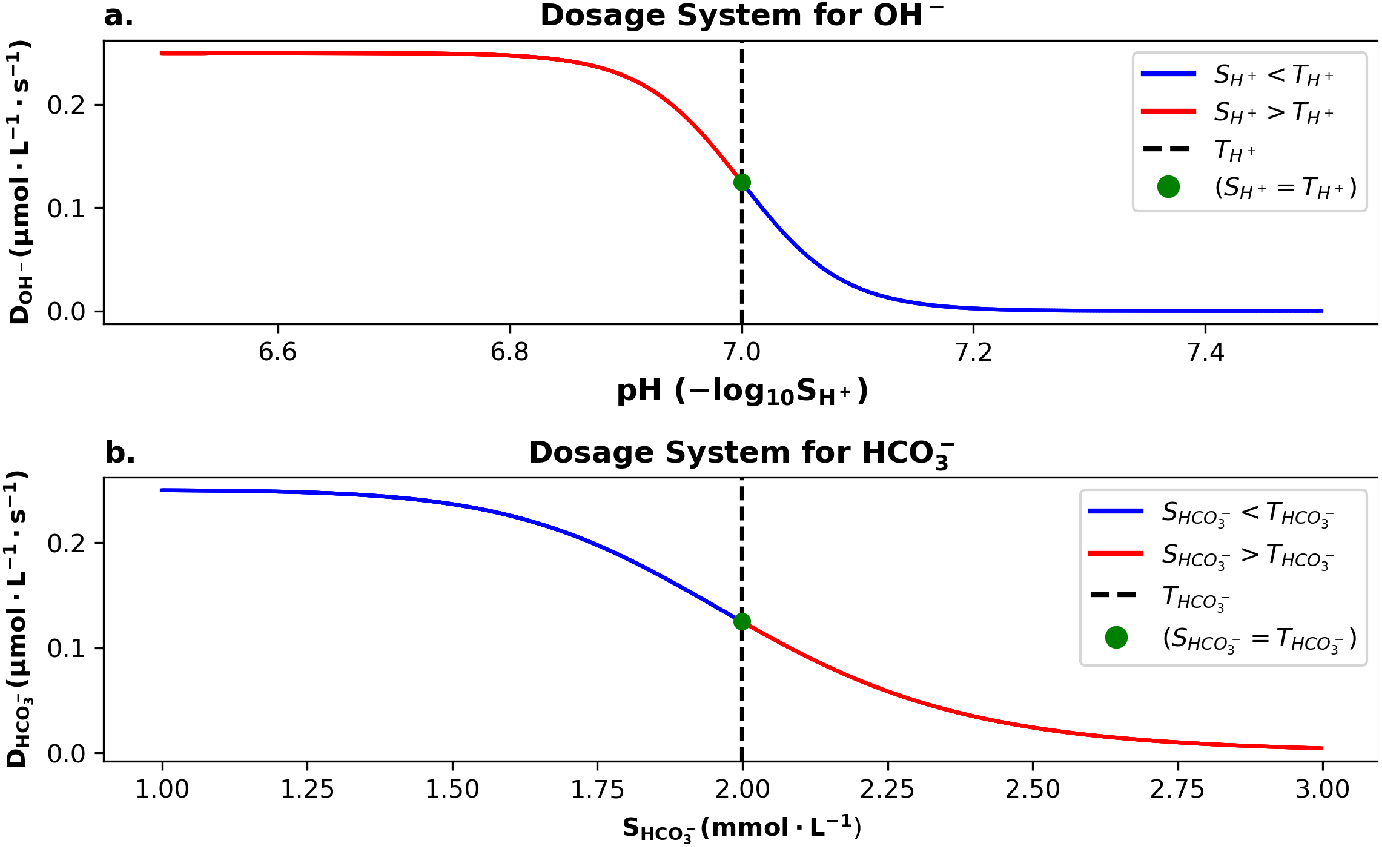
Illustration of the dosage system for *OH*^−^ and 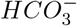 addition. Panel a) presents the dosing rate of *OH*^−^ in function of the pH. Panel b) illustrates the dosing rate of 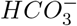 in function of the concentration of 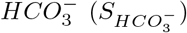 in the targeted compartment. The black dashed line indicates the threshold for the pH (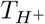 in panel a) and the targeted 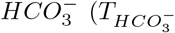 concentration in panel b). The red section of the curve indicates when the species concentration (*S*_*i*_) is above the threshold (*T*_*i*_) while the blue section shows when the species concentration is below the threshold. The green dot marks where the species concentration equals exactly the threshold.

In these dosing systems, *n* denotes the coefficient that defines the sensitivity of the system to concentrations of 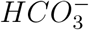 or *H*^+^. A higher value of *n* typically means a more sensitive or steep response to changes in concentration. Values of *n* used in simulations can be found in Tab. 2.

These dosing systems can be used to control either pH or alkalinity. Depending on what shall be controlled, the threshold values are computed differently. To control alkalinity, we use 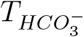 as an approximation for the alkalinity level as most of the alkalinity is present in the form of 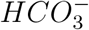 in water with pH below 8.3 [9]. For the *H*^+^ control system, 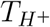 is calculated to correspond to a targeted 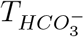 as follows [9],

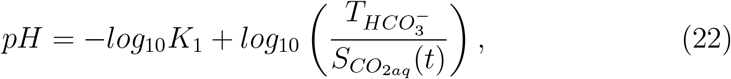

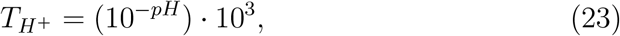

where *K*_1_ is the equilibrium constant for reaction (4) [24]. Similarly, when controlling pH with the 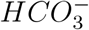 control system, we calculate 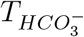 as follows:

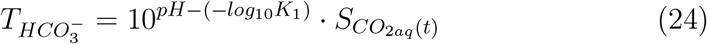

#### 2.4.3. Fish growth and excretion

Variations of water quality over time, fish growth, biomass, TAN excretion rates, and *CO*_2_ excretion rates are implemented within the model as processes occurring in the fish tank compartment. Fish growth is implemented as in [15] using the thermal growth rate coefficient approach. The growth of the fish is defined as follows,

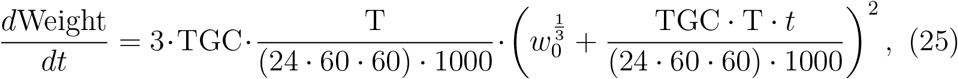

where *T* is the temperature in Celsius (°C), *TGC* is the thermal growth rate coefficient and was reported to be equal to 2.7 for fish reared between 4-14°C [25], *t* is the time in days and *w*_0_ is the initial weight in gram. Biomass can then be determined by multiplying the fish weight with the fish count. We run cycles of 14 days where the fish biomass is reduced to 50 *kg* · *m*^−3^ at the end of each cycle to investigate the effect of fish size on the water quality.

To model *CO*_2_ production over time we use a model for oxygen consumption [26, 27]. As one mole of *O*_2_ consumed will produce one mole of *CO*_2_ [25], the production of *CO*_2_ can thus be approximated by multiplying the *O*_2_ consumption by the ratio of their respective molecular weight (44*/*32). Therefore we define *CO*_2_ production rate in *mmol* · *L*^−1^ · *s*^−1^ as

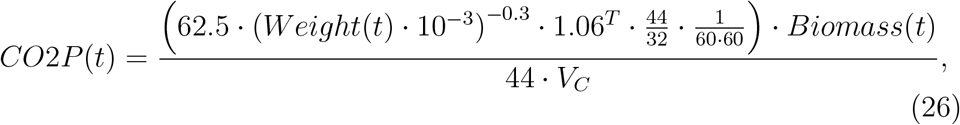

where *T* is the temperature in Celsius (°C), *Weight* is the weight of the fish in g, *Biomass*(t) is the biomass of fish in kg, *V*_*C*_ is the volume of the compartment in L. This expression takes into account the effect of temperature and body weight as small-sized fish have a higher metabolic rate than bigger fish. The model was developed in [27] using multi-regression analysis on data from measurements of oxygen consumption of a population of Atlantic salmon [26]. As different feeding regimes can induce different dynamics in the water quality parameter, the model incorporates a feeding regime of 12h intervals. To increment the impact on the *CO*_2_ excretion, we introduce an increase of 3.7% of *CO*2*P* for every hour within the feeding interval and, in contrast, a decrease of 3.7% of *CO*2*P* for every hour outside of the feeding period. The change in the production rate of for *CO*_2_ is implemented as follows,

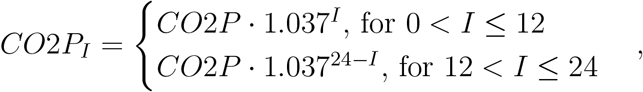

where *I* is the conversion of the given time in seconds to its equivalent fraction of one day, expressed in hours. *I* is thus a value between 0 and 24. Through this approach, our goal is to simulate a rise in fish activity during feeding times, which translates to an elevated rate of *CO*_2_ excretion. Although values for the increase of *CO*_2_ excretion rate during a 12h feeding regime have not been reported in the literature, post-prandial *O*_2_ consumption rates have been reported to increase up to 1.5-2.5 times the standard metabolic rate in salmonid [28]. We chose to simulate peak increase of about 55% of the *CO*_2_ production rate before feeding based on this information and observations of measured data for *CO*_2_. Similar to *CO*_2_ excretion, we implemented variations exerted on 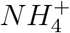 excretion induced by a 12h feeding regime. The amount of feed per day is first calculated based on the biomass present in the system. We chose to simulate a feed ratio (FR) of 2.7% body weight in accordance with the TGC. The total ammonia nitrogen in *mmol* excreted by the fish over one day is then approximated using the formula from [11],

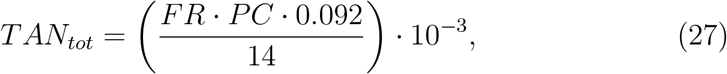

where PC is the protein content of the feed and can be found in Tab. 2. Studies indicate that peak TAN concentration occurs concurrently with peak *O*_2_ consumption rate [29]. To model peak TAN excretion over the course of a day, we fit *TAN*_*tot*_ to a beta-distribution function 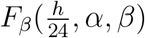 where *h* corresponds to the hour of the day (1-24) and *α* and *β* are the shape parameters of the curve. The final expression of the TAN excretion function at an instant t is given as

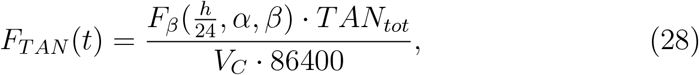

where *F*_*TAN*_ is expressed in *mmol* · *L*^−1^ · *s*^−1^ and *V*_*C*_ represents the volume of the compartment in *L* where ammonia is excreted. Parameters *α* and *β* were tested to reproduce a peak delay induced by a 12h feeding regime. The final parameters selected can be found in Tab. 2. We use a minimum threshold of 0.1 for the output of the beta distribution function so that the TAN excreted is never equal to 0. The fraction of TAN in the form of ammonia or ammonium is then calculated based on the pH of the system and the equilibrium constant of reaction (7) [30].

#### 2.4.4. Bacterial growth and nitrification process

A frequent approach for simulating bacterial growth involves employing the Monod equation. This equation describes the relation between bacterial growth rate and nutrient concentration. As in [5] and [15], we apply this strategy in dynRAS to model the growth of ammonia-oxidizing bacteria (AOB) and nitrite-oxidizing bacteria (NOB). We assume the oxygen concen-tration constant and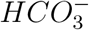, *NH*_3_, and 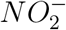 as limiting nutrients for AOB and NOB. In addition to the Monod expression, we included a term to induce a logistic growth of the bacteria due to space limitation. Equations for AOB and NOB are thus expressed as follows:

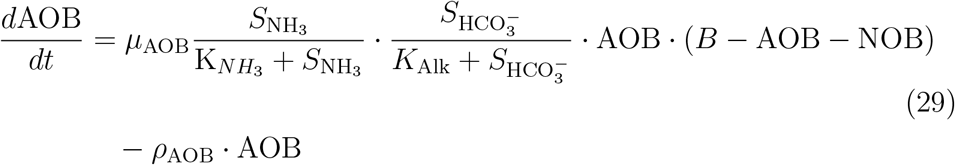

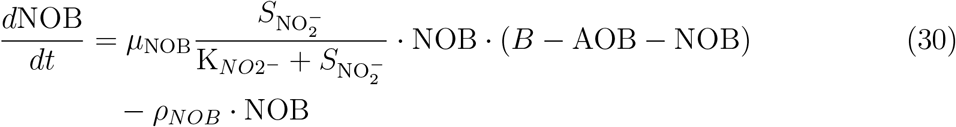

where *µ*_AOB_ and *µ*_NOB_ are the maximum growth rates for NOB and AOB, 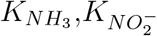 and *K*_*alk*_ are the Monod half saturation constant for *NH*_3_, 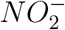 and 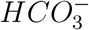, respectively. *B* represents the carrying capacity of the bio-filter for NOB and AOB in *mg* of biomass. *ρ*_*NOB*_ and *ρ*_*AOB*_ are the rate constants for the decay of the NOB and AOB.

#### 2.4.5. CO_2_ removal and Total Inorganic Carbon (TIC)

*CO*_2_ removal in the system is modelled based on the *CO*_2_ concentration in the degasser and a fixed parameter that was determined to achieve reasonable *CO*_2_ removal efficiency. The *CO*_2_ and Total inorganic carbon (TIC) removal efficiency are not variables in the model but can be determined using the model output. The TIC, *CO*_2_, and TIC removal efficiency can be calculated as follows,

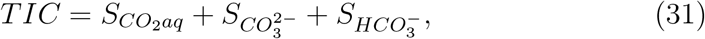

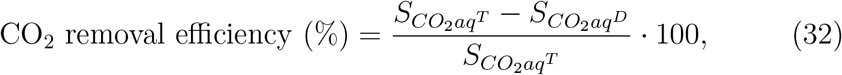

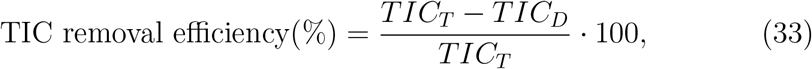

where 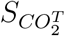 and 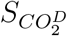 are the concentration of *CO*_2_ in the tank and the degasser, respectively. *TIC*_*T*_ represents the concentration of TIC in the tank and *TIC*_*D*_ is the concentration of TIC in the degasser.

## 3. Results

### 3.1. Model validation

We validated the model using data recorded during the experimental trial of [19] where different alkalinity levels were tested. We did not aim to have an exact reproduction of the experimental data but rather utilized the experimental results as a benchmark to validate our model. Consequently, we did not calibrate the model to fit the experimental data. We aimed to simulate a dosage system for alkalinity control, mimic the dynamic patterns induced by a 12-hour feeding regime, and reproduce existing inter-dependencies between water quality parameters.

As in the experimental design, we simulated three alkalinities 200,100 and 70 *mg*·*L*^−1^ as *CaCO*_3_ in triplicate over a period of 14 days. At the end of each of these 14-day periods fish biomass is reduced to 50 *kg* · *m*^−3^. Consistent with the original experimental design, the addition of 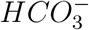 was used to control alkalinity levels around 200 *mg* · *L*^−1^ as *CaCO*_3_ while alkalinity levels were maintained at around 70 or 100 by simulating the addition of NaOH. We used an exchange rate of 75% of the total volume of the system for our simulation similar to the experimental trial. The presented data for both the simulation and experimental trial are from the fish tank compartment.

#### 3.1.1. The model qualitatively reproduces patterns observed in experimental data

The simulated dosing system effectively established the targeted alkalinity levels(Fig. 4c, Tab. 3) with minor deviations compared to the experimental data. In both simulation results and measured data, we observe that similar alkalinity can have notably different pH and *CO*_2_ values throughout a fish production cycle (Tab. 3). Mean simulated alkalinities of 101.3±9.2 *mg* · *L*^−1^ and 102.4 ± 8.7 *mg* · *L*^−1^ as *CaCO*_3_ demonstrate a pH of 6.97 ± 0.11 and 7.20 ± 0.14 respectively, with a mean *CO*_2_ of 12.3 ± 2.4 *mg* · *L*^−1^ and 7.3 ± 1.9 *mg* · *L*^−1^.

**Fig. 4.**
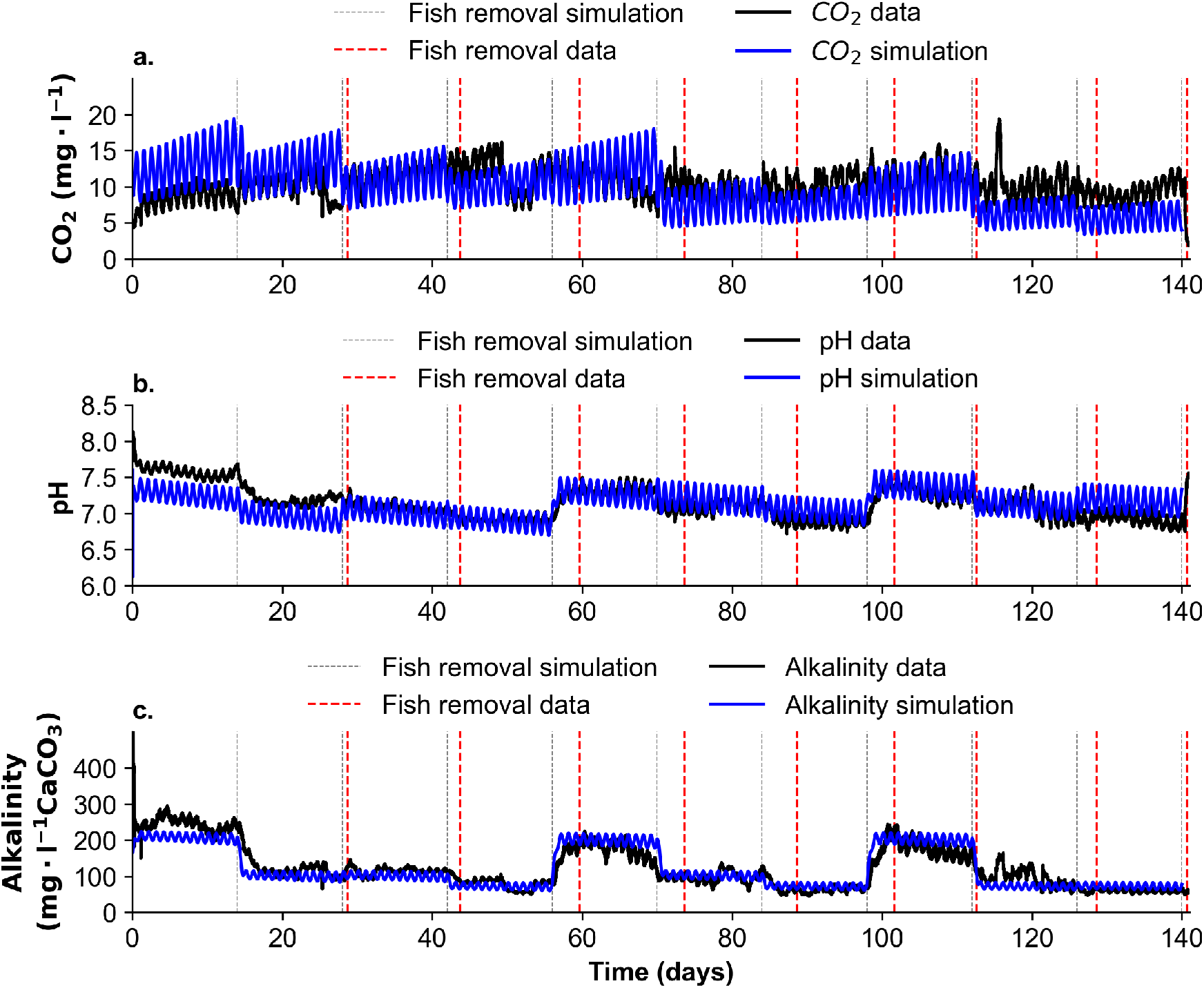
Comparison of simulation results and experimental data. Experimental data are represented by black lines and blue lines represent the simulation results. Panel a) presents results for *CO*_2_ concentration (*mg* · *L*^−1^) over 140 days. Panel b) presents results for pH and panel c) for alkalinity levels (*mg* · *L*^−1^ as *CaCO*_3_).

**Table 3:** Mean alkalinity, pH and *CO*_2_ followed by the standard deviation for both simulation results and measured data. The mean is calculated over a period of 10 days within the treatment period of 14 days, excluding days 1 to 3 which are used to set up the treatment in the experimental trial and day 14, which also corresponds to the first day of the subsequent treatment.

| Period<br>(days) | Targeted Alkalinity<br>($\text{mg} \cdot \text{l}^{-1}$ as $\text{CaCO}_3$) | Alkalinity ($\text{mg} \cdot \text{l}^{-1}$ as $\text{CaCO}_3$) | | pH | | $\text{CO}_2$ ($\text{mg} \cdot \text{l}^{-1}$) | |
| --- | --- | --- | --- | --- | --- | --- | --- |
|  |  | Simulation | Data | Simulation | Data | Simulation | Data |
| 3-13 | 200 | $207.7 \pm 9.9$ | $245.5 \pm 19.4$ | $7.27 \pm 0.12$ | $7.55 \pm 0.07$ | $12.7 \pm 3.0$ | $8.8 \pm 1.2$ |
| 17-27 | 100 | $101.3 \pm 9.2$ | $114.2 \pm 11.1$ | $6.97 \pm 0.11$ | $7.16 \pm 0.05$ | $12.3 \pm 2.4$ | $10.2 \pm 1.5$ |
| 31-41 | 100 | $101.1 \pm 9.1$ | $113.1 \pm 8.7$ | $7.02 \pm 0.11$ | $7.08 \pm 0.05$ | $10.8 \pm 2.2$ | $11.9 \pm 1.3$ |
| 45-55 | 70 | $71.9 \pm 7.5$ | $75.2 \pm 14.5$ | $6.90 \pm 0.11$ | $6.92 \pm 0.04$ | $10.1 \pm 1.7$ | $11.6 \pm 2.6$ |
| 59-69 | 200 | $201.9 \pm 11.8$ | $177.3 \pm 19.5$ | $7.29 \pm 0.13$ | $7.33 \pm 0.07$ | $11.7 \pm 2.9$ | $10.7 \pm 1.8$ |
| 73-83 | 100 | $102.4 \pm 8.7$ | $95.3 \pm 16.9$ | $7.20 \pm 0.14$ | $7.10 \pm 0.09$ | $7.3 \pm 1.9$ | $9.6 \pm 1.5$ |
| 87-97 | 70 | $72.0 \pm 7.1$ | $60.7 \pm 7.1$ | $7.04 \pm 0.11$ | $6.87 \pm 0.06$ | $7.4 \pm 1.5$ | $10.4 \pm 1.5$ |
| 101-111 | 200 | $203.8 \pm 11.4$ | $176.4 \pm 22.7$ | $7.39 \pm 0.14$ | $7.29 \pm 0.11$ | $9.4 \pm 2.5$ | $11.5 \pm 1.9$ |
| 115-125 | 70 | $73.3 \pm 7.4$ | $97.6 \pm 24.2$ | $7.13 \pm 0.13$ | $7.08 \pm 0.13$ | $6.2 \pm 1.5$ | $10.3 \pm 2.5$ |
| 129-139 | 70 | $72.6 \pm 7.0$ | $58.9 \pm 3.7$ | $7.18 \pm 0.13$ | $6.92 \pm 0.08$ | $5.5 \pm 1.4$ | $9.2 \pm 1.3$ |

The model results were comparable to the experimental data in terms of the daily patterns introduced by feeding observed for *CO*_2_, pH, and TAN (Figs. 4, 5, 6). The observed daily peaks in *CO*_2_ in the RASlab experiments were accurately captured in the simulations occurring toward the end of the feeding period (Figs. 4a, 5a). The daily fluctuations of TAN concentration are also well depicted by the model as shown in Fig. 6. Unfortunately, due to limited access to the facility, TAN concentration could not be assessed from 21:00 to 7:00, therefore the maximum TAN concentration over the day could not be experimentally asserted. Nonetheless, the trends observed in both the modelled and empirical data (Fig. 6) suggest that the peak TAN level is likely to occur at the end of the feeding period. In addition, as the model takes into account fish growth and fish removal, we observe a general decrease in *CO*_2_ concentration for the simulated result that is also depicted in the measurement data (Fig. 4a). However, the drop of *CO*_2_ concentration at each fish removal event (every 14 days) is more marked in the simulated data as the fish are removed instantaneously at regular intervals as opposed to the experiment where fish were removed at slightly irregular intervals.

**Fig. 5.**
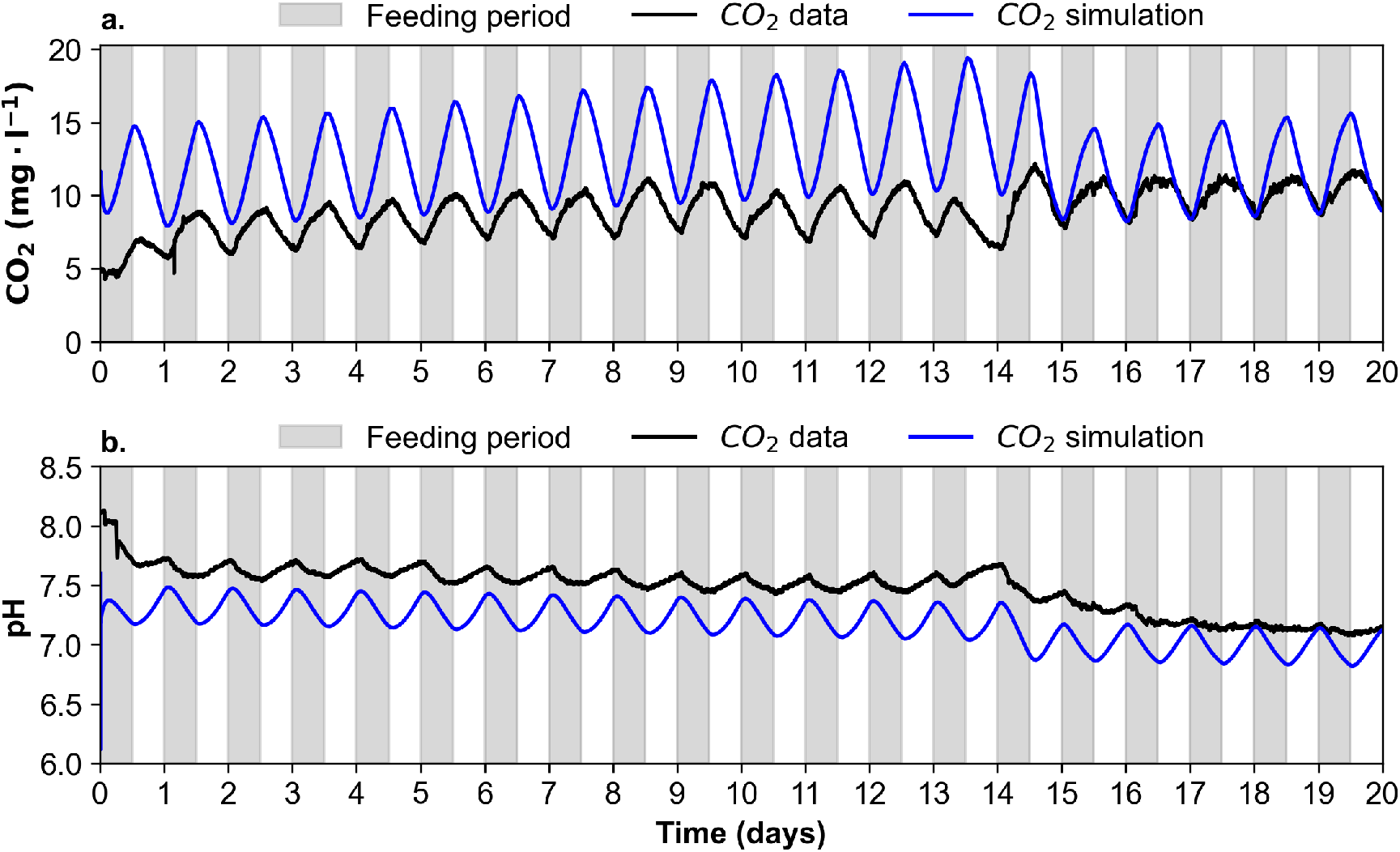
Zoomed-in view of the first 20 days of both the simulation results and the experimental data. The feeding periods are highlighted with grey background bars. Panel a) presents the *CO*_2_ concentration and panel b) presents results for pH. For both panels, black lines represent the experimental data and blue lines indicate the simulation results.

**Fig. 6.**
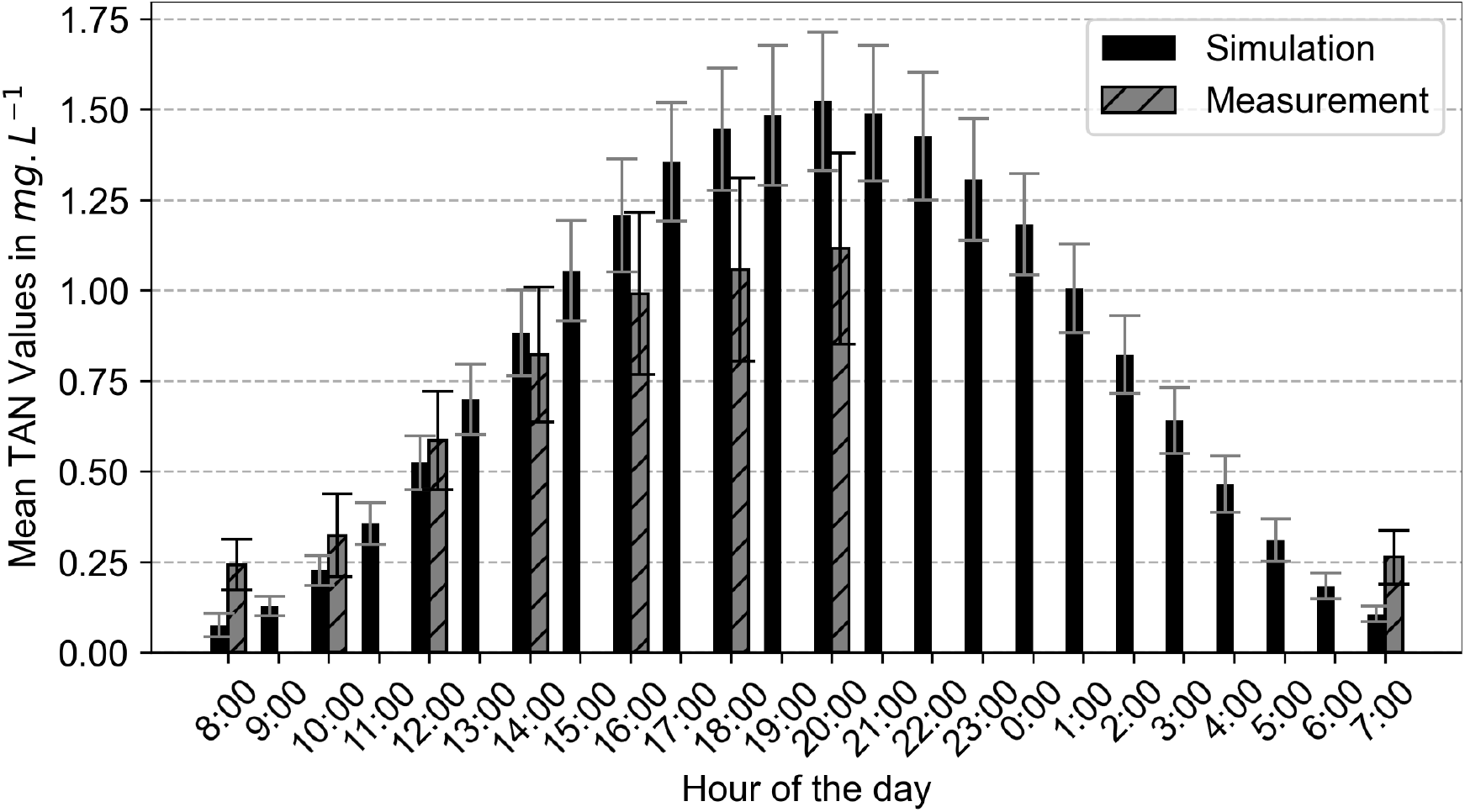
Diurnal Variation of Total Ammonia Nitrogen (TAN) Levels in the Tank: This figure illustrates the mean TAN concentration, along with its standard deviation, observed over a 24-hour period. Black bars indicate TAN concentrations from the simulation, while grey hatched bars depict actual measurements obtained from the tank. Data collection was conducted every two weeks, with samples taken every 2 hours throughout the feeding period, which extends from 8:00 to 20:00. An extra sampling was taken just prior to the start of the next feeding period to capture TAN levels before feeding resumed.

#### 3.1.2. Inter-dependencies between pH, CO_2_ and alkalinity are well captured

While qualitatively accurate, the model overestimates the *CO*_2_ production at the beginning of the simulation and underestimates the production towards the last days of the simulation. We also observe in general larger amplitudes for simulated *CO*_2_ over the day compared to the measurement data (Fig. 4a). Higher mean simulated *CO*_2_ levels can be observed for targeted alkalinity 200 which are not observed in the experimental data. This observation is also correlated with a lower *CO*_2_ and TIC removal efficiency at targeted alkalinity 200 (Tab. 4). Because continuous measurements for the outlet of the *CO*_2_ stripper were not recorded during the experimental trial, model results could not directly be compared with actual data. However, significantly higher *CO*_2_ removal efficiency for nominal alkalinity 100 and 70 were also found in [19] as well as a lower TIC removal efficiency for alkalinity 200.

**Table 4:**
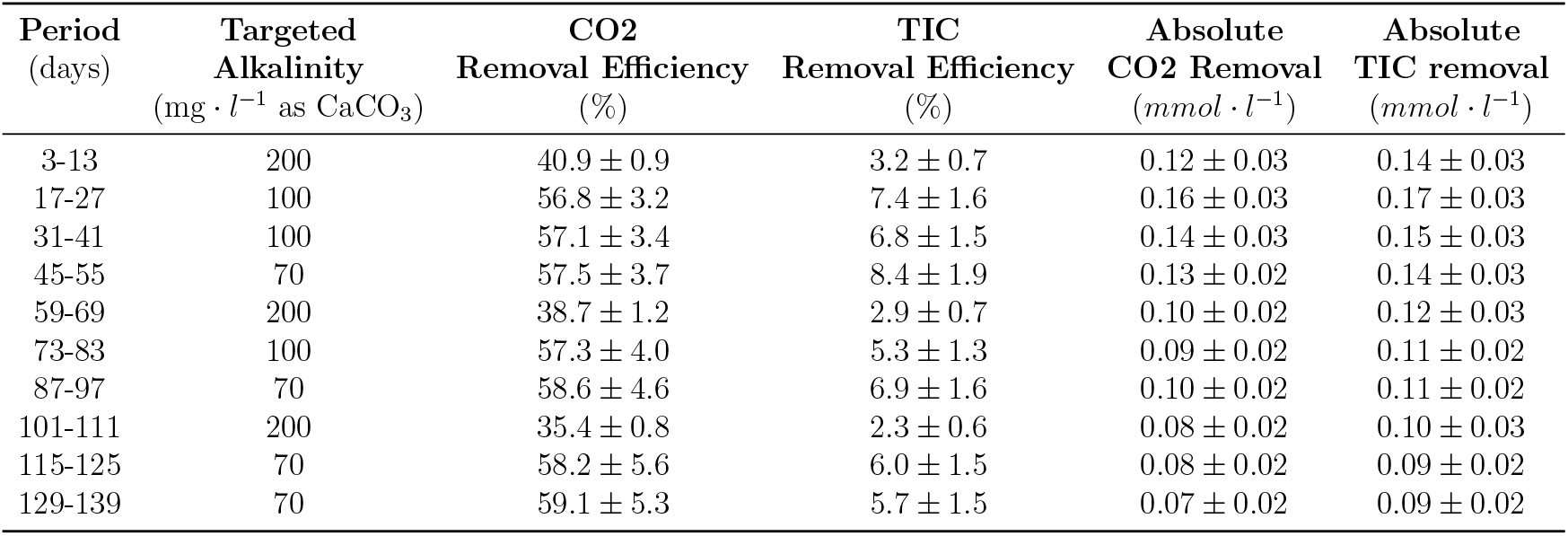
Simulation result for mean removal efficiency for *CO*_2_ and TIC and their absolute value in *mmol* · *L*^−1^

Despite these discrepancies, the model effectively captures the inverse relationship between *CO*_2_ and pH, with the lowest pH coinciding with the daily peak in *CO*_2_ levels, consistent with the experimental data depicted in Fig. 5. In both the simulation and the measured, data higher nominal alkalinity of 200 *mg* · *L*^−1^ resulted in elevated pH levels compared to lower alkalinity (Fig. 4, Tab. 3).

### 3.2. Exploring alkalinity level and supplement strategies using the dynRAS model

As the model has proven its capability to reflect the existing interdependencies within the system, we used the model to simulate 3 different scenarios:

- **Scenario 1**: Only bicarbonate 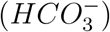 is used to establish alkalinity levels of 200 and 70.
- **Scenario 2**: Alkalinity levels of 200 and 70 are maintained exclusively using the NaOH dosing system.
- **Scenario 3**: We combined the two dosing systems to control the same alkalinity levels. In this last scenario, we modify the Hill function of each dosage system to take into account the *CO*_2_ level in the system.

Each scenario is simulated for 140 days, with fish being removed every 14 days to maintain the fish biomass at 50 *kg* · *m*^−3^. Through the simulations of these scenarios, we aim to demonstrate how the model can be used to investigate the optimal alkalinity level as well as the optimal buffer or base supplement strategy.

#### 3.2.1. Scenario 1 and 2: Comparison of base or buffer addition on the system dynamic

Results from scenarios 1 and 2 illustrate the trade-offs associated with each supplement. Higher *CO*_2_ levels are observed in the system regardless of the alkalinity level when 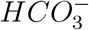 is used as a supplement (Fig. 7). Notably, we observe that *CO*_2_ levels can reach concentrations above 20 *mg* · *L*^−1^ when 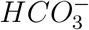 supplements are used within day 14 to day 42 of the simulation (Fig. 7a). In addition, the efficiency of *CO*_2_ removal and therefore the efficiency of TIC removal is diminished due to the introduction of inorganic carbon when using 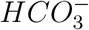 compared to the use of *NaOH* (Fig. 8).

**Fig. 7.**
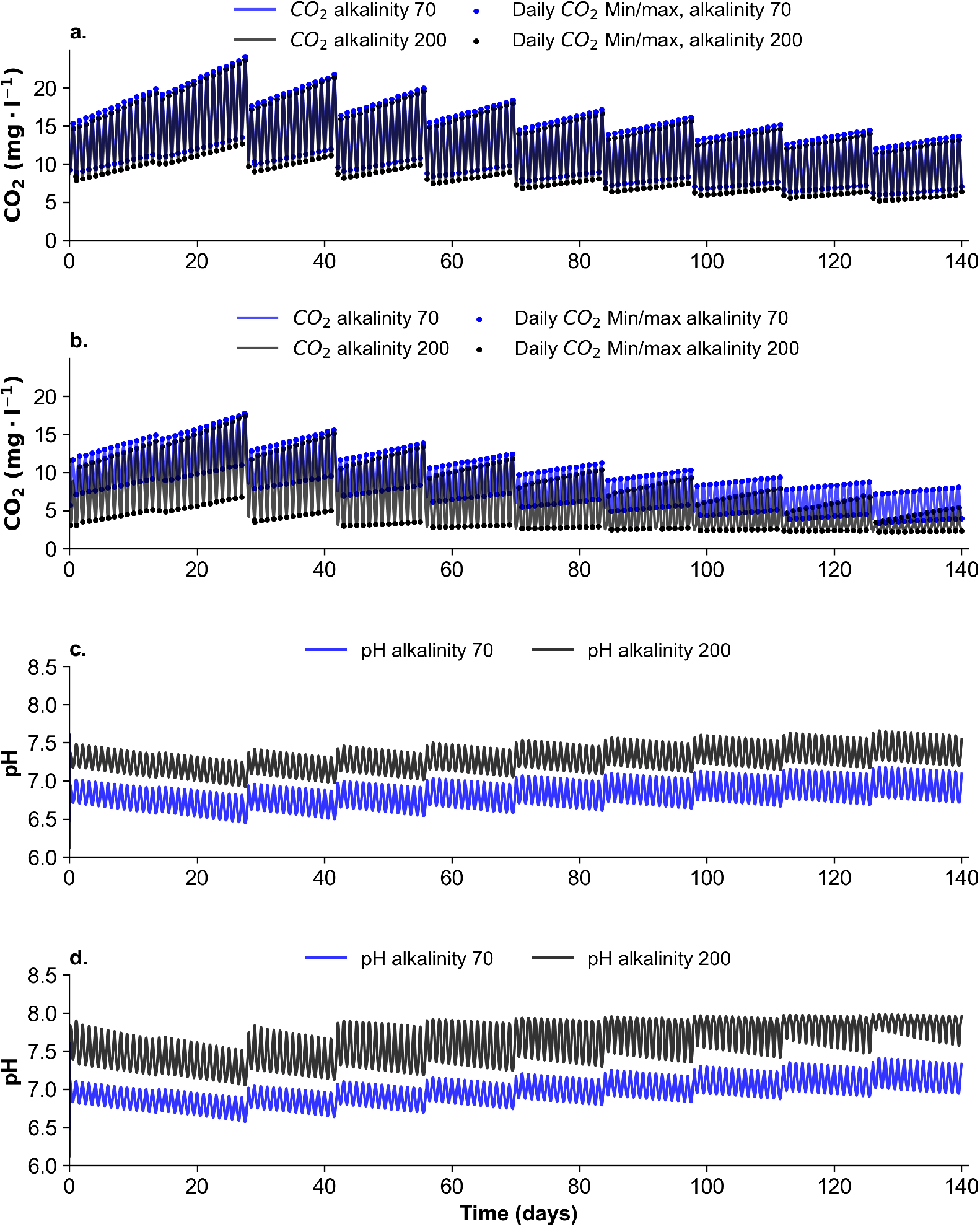
Simulation results for *CO*_2_ concentrations and pH levels for scenario 1 (panels a and c) and scenario 2 (panels b and d). Blue lines represent simulation results for targeted alkalinity 70 *mg* · *L*^−1^ as *CaCO*_3_ and black lines represent simulation results for alkalinity 200 *mg* · *L*^−1^ as *CaCO*_3_. Black and blue dots were added for better visibility of the daily maximum and minimum *CO*_2_ levels in the system.

**Fig. 8.**
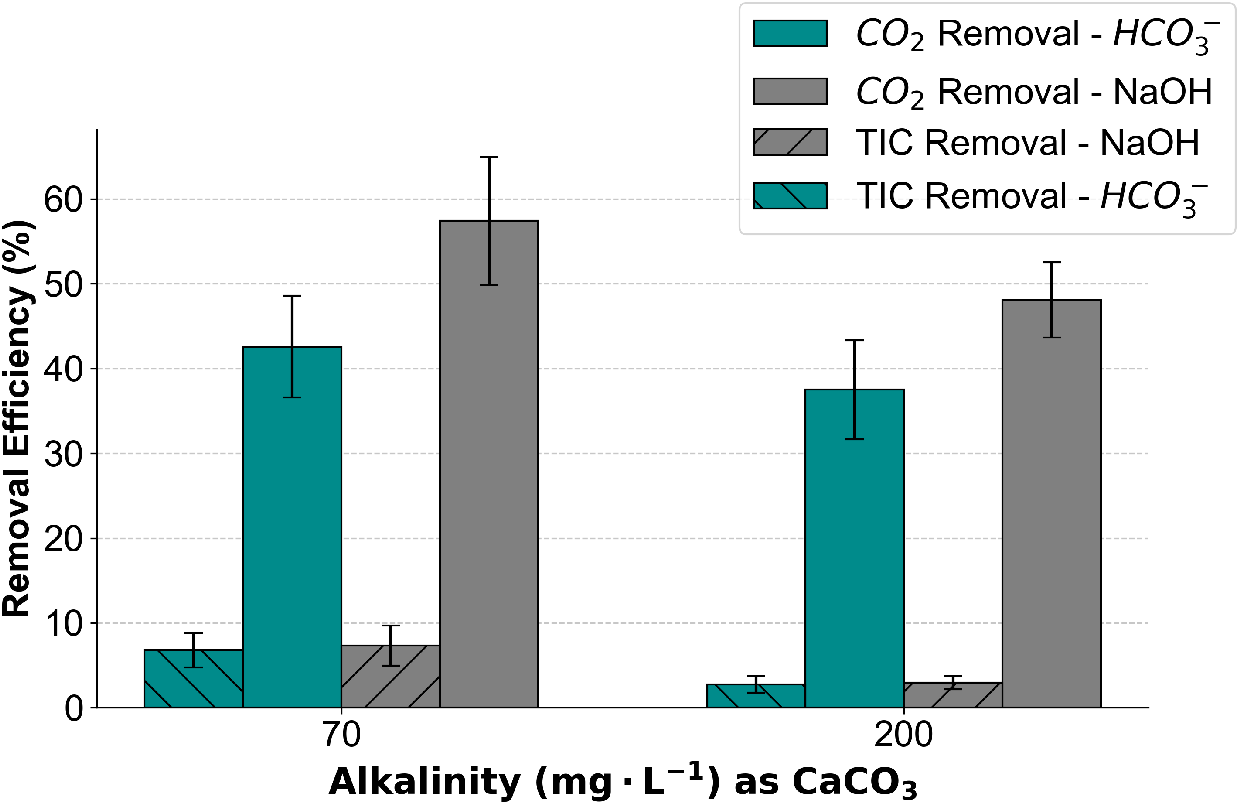
Simulation result for *CO*_2_ removal and TIC removal for alkalinity 70 and 200 *mg* · *L*^−1^ including the standard deviation. Cyan bars represent scenario 1 and grey bars represent scenario 2.

While the use of *NaOH* allows for operating the system at lower *CO*_2_ levels and higher pH (Fig. 7), higher pH leads to higher *NH*_3_ levels despite the lower TAN concentration in the system (Fig. 9c,d). Particularly, *NaOH* controls alkalinity by increasing pH and shifting reaction (2) toward 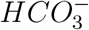.

**Fig. 9.**
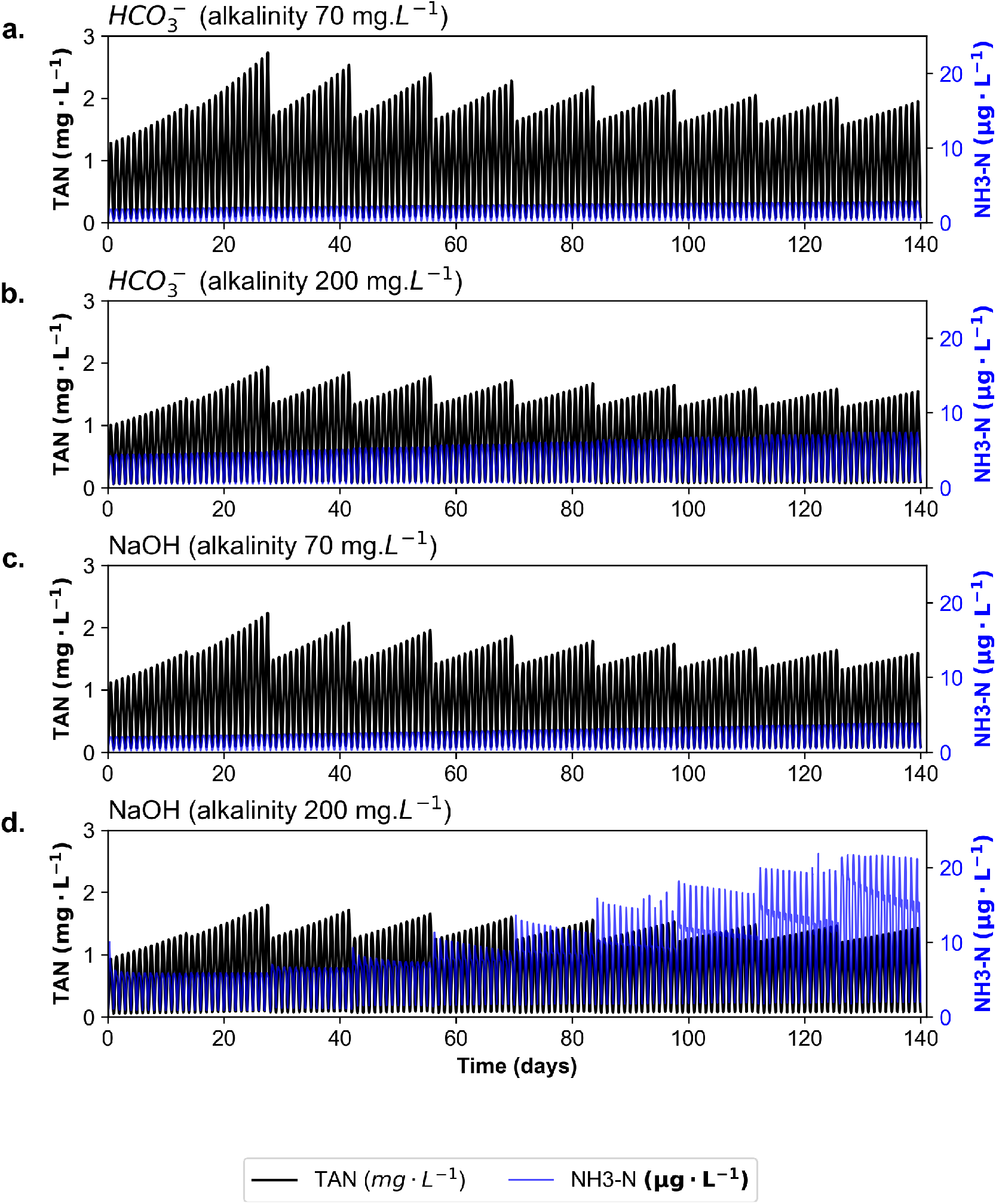
Simulation results for TAN and *NH*_3_-N for alkalinity 70 and 200 *mg* · *L*^−1^ as *CaCO*_3_ for scenario 1 (panel a and b). and scenario 2 (panel c and d). Blue lines represent the *NH*_3_-N concentration and black lines the TAN concentration in the fish tank.

As no inorganic carbon is added to the system, we observe that the required pH to maintain an alkalinity level of 200 increases as the *CO*_2_ production decreases when using *NaOH*. This leads to situations where the fraction of ammonia reaches the maximum recommended concentration (Fig. 9d) [10].

Overall, the results show that the choice between using a base or a buffer supplement is highly dependent on the *CO*_2_ production in the system. Using *NaOH* when *CO*_2_ production is low will lead to too high pH. Conversely, using 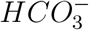 when *CO*_2_ production is high would lead to potentially harmful *CO*_2_ levels for the fish. Therefore, in scenario 3 we explore the implementation of a dosage system combining the addition of both 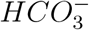 and *NaOH* based on the *CO*_2_ concentration in the system.

#### 3.2.2. Scenario 3: Dosing with both 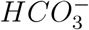 and NaOH

Instead of using either *NaOH* or 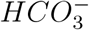 to control pH and alkalinity, using a combination of these compounds could prove to be more beneficial to the system. The addition of NaOH is more advantageous in scenarios with high *CO*_2_ levels while the opposite is true for 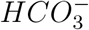. To explore this further through simulations, we modified the Hill function for the dosage systems to account for *CO*_2_ concentration as follows,

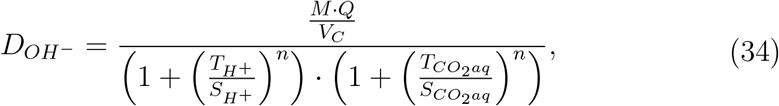

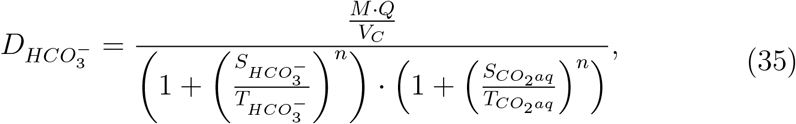

where 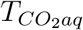 is the threshold concentration used to modulate the addition of *NaOH* and 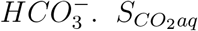 is the *CO*_2_ concentration in the compartment. We chose a threshold of 10 *mg* · *L*^−1^. If the *CO*_2_ concentration is below that threshold the addition of *NaOH* is decreased while the addition of 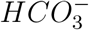 increases. Conversely, if the *CO*_2_ concentration is above the threshold then the dosing of *NaOH* increases while the dosing of 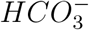 decreases. This approach allows us to control alkalinity and optimize the use of the base and buffer supplement in accordance with the *CO*_2_ concentration in the system.

As the *CO*_2_ decreases with time (Fig. 10a) we observe a decrease in the dosing rate of *NaOH* while the dosing rate of 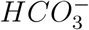 increases (Fig. 11). Differences in the dosing rate for both supplements can be observed depending on the targeted alkalinity level. An alkalinity level of 200 leads to higher operating pH, and thus slightly lower *CO*_2_ concentration in the system (Fig. 10) compared to alkalinity 70. Thereby, the dosing rate for 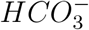 is higher for targeted alkalinity 200 than for alkalinity 70 (Fig. 11). Conversely, the dosing rate of *NaOH* is higher for alkalinity 70 compared to its dosing rate at alkalinity 200 (Fig. 11). Using a dosage system that adapts the addition of both supplements following the *CO*_2_ levels mitigates situations with high *CO*_2_ concentration or high pH that could occur if only one supplement is used. In fact, *CO*_2_ levels from scenario 3 (Fig. 10a) are lower from day 0 to day 42 of the simulation compared to the *CO*_2_ level over the same days when only 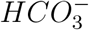 is used in scenario 1 (Fig. 7a). In addition, lower pH values are observed from day 42 to day 140 of the simulation in scenario 3 (Fig. 10b) compared to scenario 2 where only *NaOH* is added (Fig. 7d). These lower pH values allow operating the system with lower and safer *NH*_3_ − *N* levels compared to scenario 2, particularly when an alkalinity level of 200 *mg* · *L*^−1^ is targeted (Figs. 12a and 9d).

**Fig. 10.**
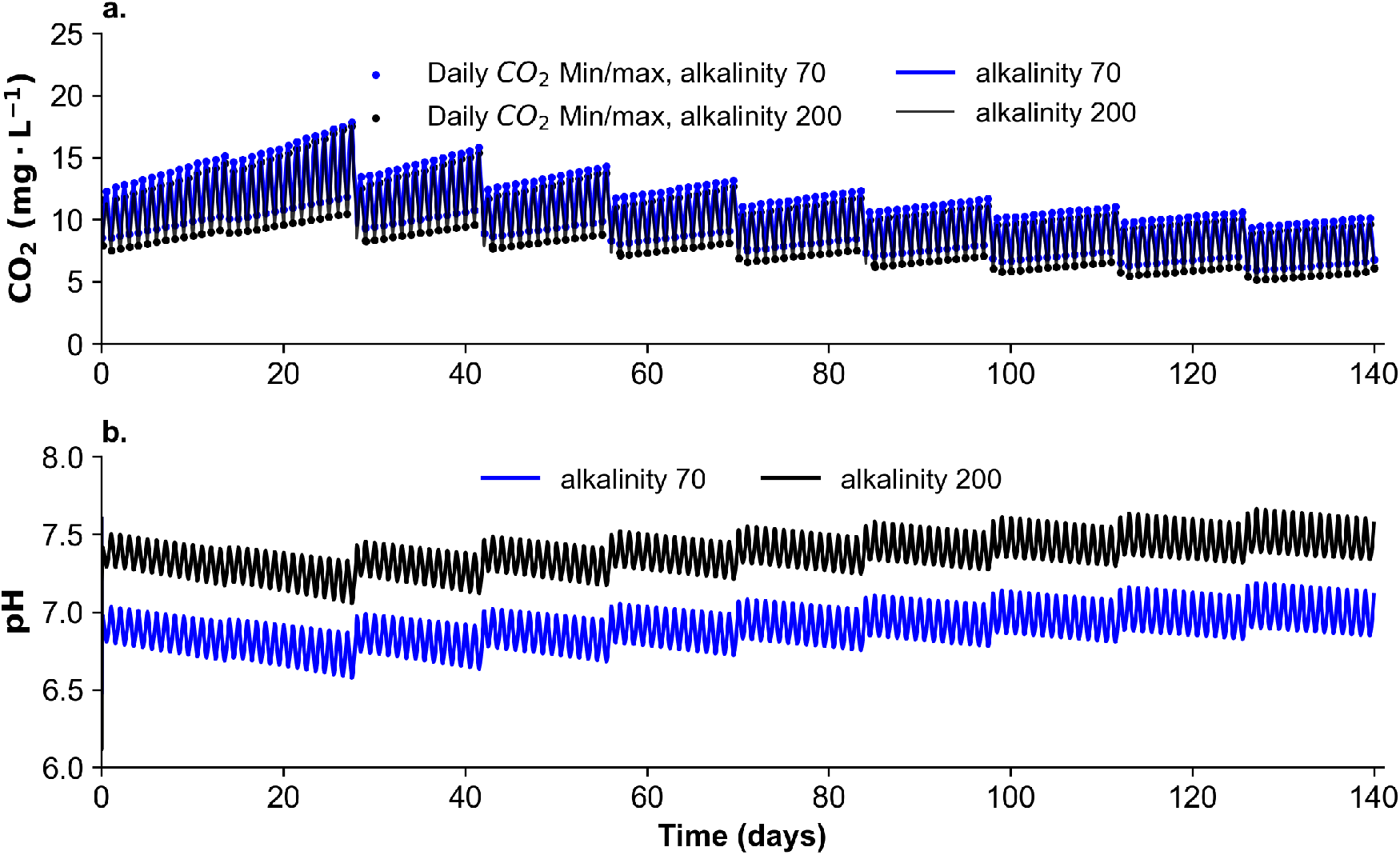
Simulation results for scenario 3 for a) *CO*_2_ and b) pH. Blue lines represent simulation results obtained for targeted alkalinity of 70 *mg* ·*L*^−1^ and black lines correspond to simulation results obtained for alkalinity 200 *mg* · *L*^−1^ as *CaCO*_3_.

**Fig. 11.**
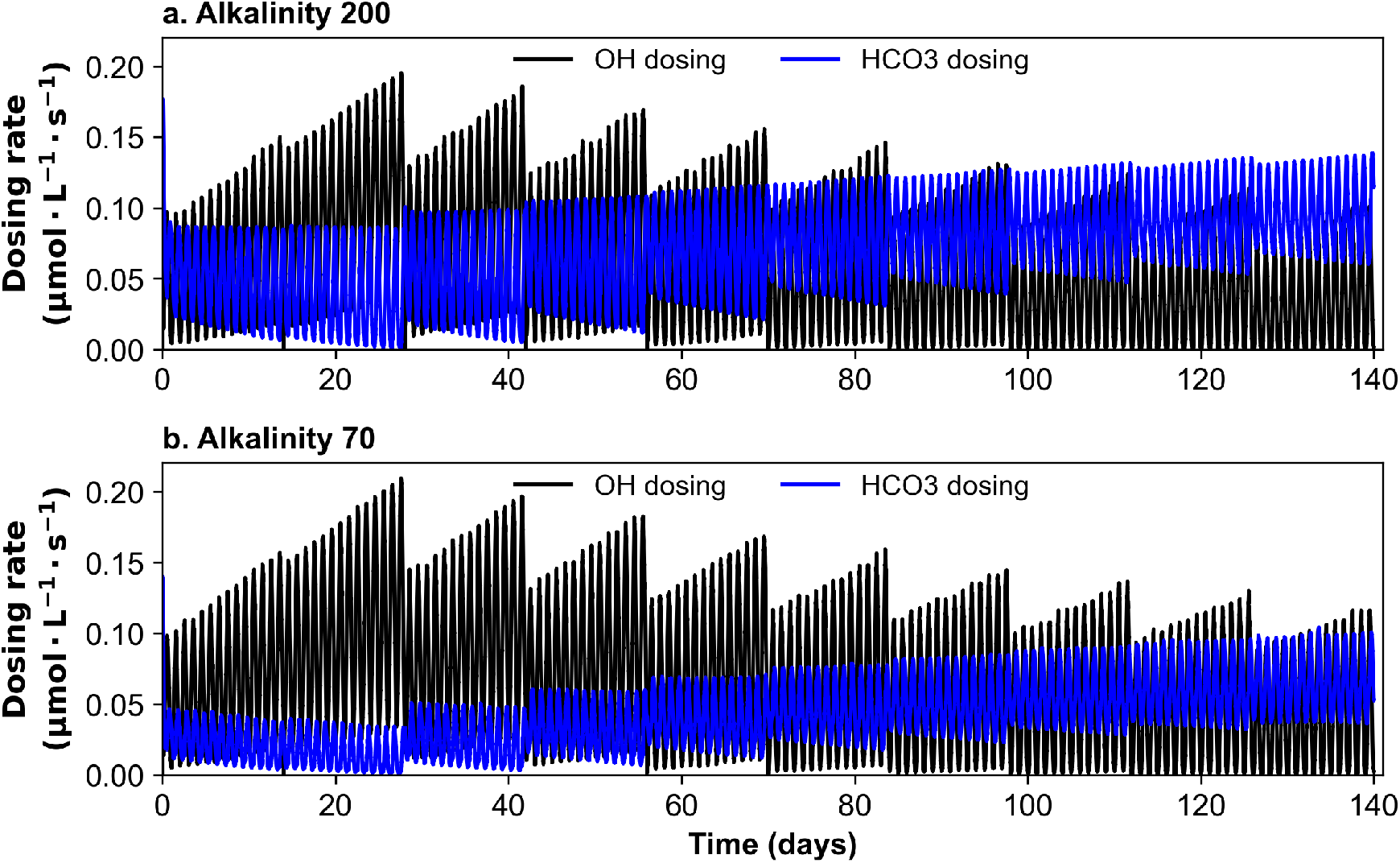
Dosing rates of *NaOH* and 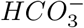 for a) alkalinity 200 and b) alkalinity 70. Blue lines correspond to the dosing rate of 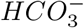 while black lines correspond to the dosing rate of *NaOH*.

**Fig. 12.**
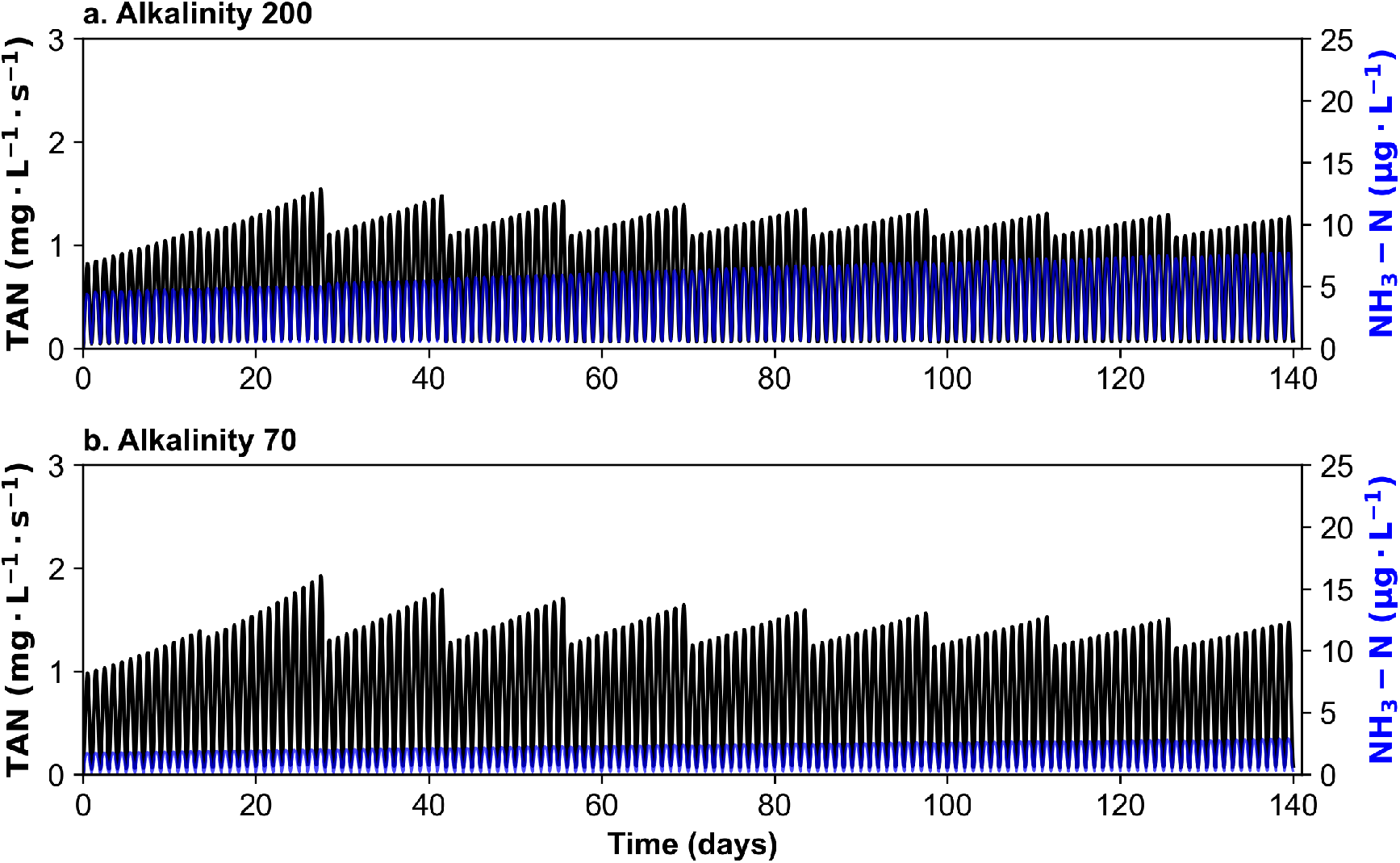
Simulation results for scenario 3 for TAN and *NH*_3_-N concentration. Results for targeted alkalinity 200 *mg* · *L*^−1^ as *CaCO*_3_ are represented in panel a) and results for alkalinity 70 *mg*·*L*^−1^ as *CaCO*_3_ are represented in panel b). Blue lines represent simulation results obtained for *NH*_3_-N concentration and black lines correspond to simulation results obtained for TAN concentration

Overall these results show that alternating the use of *NaOH* and 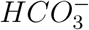 based on the *CO*_2_ concentration appears to be the most beneficial strategy for the modelled system in terms of *CO*_2_, pH and *NH*_3_ levels.

#### 3.2.3. Scenarios 1, 2 and 3: Comparing the effect of two different alkalinity levels on the system dynamic

Results from all scenarios indicate that independent of the dosing strategy used, a higher alkalinity of 200 will induce higher operational pH than when setting an alkalinity level of 70 (Fig. 7b, Figs. 10c,d). Running the system at high alkalinity and therefore high pH implies that a more significant fraction of ammonia can be present in the system despite having lower TAN overall (Figs. 9b,d, Fig. 12c) compared to running the system at lower alkalinity (Figs. 9a,c, Fig. 12d). Particularly, the pH increases over time with the decrease of *CO*_2_ production. This increase is also accentuated by the control system to maintain a stable alkalinity level throughout the simulation. This will lead to situations where maintaining an alkalinity of 200 when the *CO*_2_ production is at its lowest becomes challenging and may be detrimental to the system due to the higher fractions of *NH*_3_ − *N*. Such scenarios are observable from day 98 to day 140 of the simulations (Figs. 9b,c, Fig. 12c). Additionally, *CO*_2_ removal efficiency was observed to decrease at an alkalinity of 200 compared to 70. This is particularly marked when using solely *NaOH* (Fig. 8).

Conversely, while setting an alkalinity of 200 leads to high pH, setting an alkalinity of 70 at the beginning of the simulation (days 0-28) when the *CO*_2_ production is at the highest leads to low pH (Figs. 7c,d, Fig. 10b). These low pH can be sub-optimal for the bio-filter performance. In this scenario, setting an alkalinity of 200 from day 0 to day 28 of the simulation appears more beneficial for the system as it establishes higher pH as well as slightly lower *CO*_2_ (Fig. 7, Fig. 10).

Maintaining a constant alkalinity level throughout the production cycle may not be the most effective strategy due to the system’s dynamic nature and the implications of setting one specific alkalinity level. The results suggest that the optimal alkalinity level and choice of base or buffer supple-mentation should be assessed based on the *CO*_2_ production in the system.

## 4. Discussion

The notion of optimal in RAS is not immutable when considering water quality management. Because these systems include feedback mechanisms between fish growth, waste production, and water quality, what is considered optimal must be adaptable to maintain system balance. This makes designing practices for water quality control in RAS challenging. When fish growth and feeding regimes introduce long-term and short-term variations in metabolite levels, adding chemicals to control pH and alkalinity can significantly impact the toxicity of these metabolites if not carefully designed. Using mathematical modelling to predict these variations and understand existing interdependencies in the system is a powerful approach to refining practices in water quality management.

### 4.1. Model discrepancies compared to measurement data

Although the model qualitatively reproduced the experimental data, some disparities were observed. Differences in *CO*_2_ levels are observed at the beginning and end of the simulation compared to measured data. The differences observed at the start of the simulation potentially originate from the model used for *CO*_2_ excretion. The model used is a model for *O*_2_ consumption and was extracted from data for fish between 0.200 kg to 3.3 kg fish [27]. This may explain the overestimation for *CO*_2_ excretion at the beginning of the simulation as the fish approach 200g around day 40. However, we could not find a suitable model that would cover the full range of fish weight in our simulation.

In addition, the fish in the model are in “perfect” growth conditions; they grow at a constant rate based on their initial weight whereas environmental disturbance occurs in real-life conditions. A slightly lower weight was measured in the observed data towards the end of the experiment compared to the simulation (fish growth data are available in the supplementary material Fig. A.13). As the *O*_2_ consumption rate decreases with the increase in the weight [26], the *CO*_2_ excretion rate will follow the same trend. At the same biomass, bigger fish will excrete less *CO*_2_ than small fish.

Larger variations for pH and *CO*_2_ are observed in the simulation compared to the experimental data, as reflected by the larger standard deviation in Tab. 3. In the simulation, alkalinity is more tightly controlled than pH which could explain these larger daily variations. We use the model to run an additional scenario to illustrate how alkalinity will change when pH is maintained and *CO*_2_ production changes (Supplementary material, Fig. A.14). When *CO*_2_ increases, a fraction is converted to 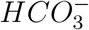, and this fraction is pH-dependent. If pH is maintained at a certain level, more *CO*_2_ will convert to 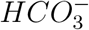 than if pH drops reactions (4) and (5)). As *CO*_2_ increases, when alkalinity is controlled, the system adjusts the dosage of base or buffer based on 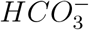 concentration, reducing additions if 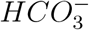 is above the thresh-old. This results in larger pH drops and larger simulated *CO*_2_ variations compared to experimental data. Discrepancies between the model simulation and empirical data for *CO*_2_ levels could be attributed to the chosen parameters, which were determined for a salinity of 35 ppt, while the pilot scale salinity was 17 ppt. Typically, higher salinity leads to lower *CO*_2_ solubility [22], resulting in reduced *CO*_2_ concentration in the system. This is converse with the observed results, suggesting that the model discrepancies might be due to other influencing factors.

The model results also present higher *CO*_2_ concentrations at alkalinity 200 compared to alkalinity 70 and 100 which is not featured in the data. In the simulation, the model strictly uses 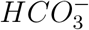 to maintain alkalinity at 200 and strictly uses *NaOH* to control alkalinity at 70 and 100 with no additional inorganic carbon source apart from the water exchanges. The higher *CO*_2_ levels at alkalinity 200 are thus likely due to the dosing of 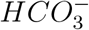 in the system. This is supported by scenario 1, which shows generally higher *CO*_2_ levels when dosing with 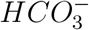 (Fig. 7a).

While there were discrepancies with the experimental dataset, the primary objective of this model was to replicate the relation between pH, alkalinity and *CO*_2_ as well as the daily trends of *CO*_2_ excretion and TAN under a 12h feeding regime. In regard to the overall simulation results, this objective was achieved.

### 4.2. The dynRAS model: a potential tool to understand the dynamic of the system and refine water management practices

Although a wide range of alkalinity levels are reported in the literature, when comparing different alkalinities, authors often try to settle over one particular level. In the studies conducted by Jafari et al., (2024) [19] and Summerfelt et al., (2015) [33], various alkalinity treatments were set in triplicates of 14 days each. While these studies provided valuable insights, long-term changes in system balance, particularly, long-term variations in *CO*_2_ production may have been overlooked with these settings. However, extending the assessment period to capture long-term effects would be prohibitively costly and necessitate running multiple RAS in parallel. In this context, model simulations become valuable for understanding long-term system dynamics at a lower cost. While consistent with findings from [19], our model simulations offer more nuanced insights into optimal alkalinity levels, offering a deeper understanding of when these levels can be the most effective. Results showed that setting an alkalinity level of 200 toward the end of the production cycle in scenarios where the biomass is maintained relatively constant, can result in higher pH levels with potentially higher ammonia levels (Figs. 7c,d, and Figs. 12b,d). Although, the upper recommended limit of 0.0125 *mg* · *L*^−1^ [10, 11] for ammonia may not be reached when using 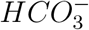 as dosing sup-plement, targeting lower alkalinity toward the end of the production cycle would reduce the use of dosing agents (Fig. 11) while keeping lower levels of ammonia (Fig. 12) and acceptable *CO*_2_ levels in the system (Fig. 10). However, while setting lower alkalinity towards the end of the production cycle can be more beneficial, setting higher alkalinity levels at the start of the production may be needed to maintain pH and *CO*_2_ within a suitable range for salmonids. Additionally, if the *CO*_2_ production is high and pH is maintained, it is likely that the alkalinity in the system will also be high as more inorganic carbon will be in the form of 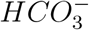 (Supplementary material, Fig. A.14).

Understanding how the system balance changes over time is important for selecting the appropriate alkalinity level but also for efficient control. Results from scenarios 1, 2 and 3 underline the significance of selecting the appropriate supplement at the optimal time in the production cycle. While using *NaOH* to control the pH and, by extent, the alkalinity level will reduce *CO*_2_ concentration, it may not be the optimal supplement at every stage of the production cycle. When using *NaOH* to regulate pH and maintain alkalinity, fish biomass, fish weight and thereby *CO*_2_ production have to be closely considered. The *CO*_2_ production from the fish must be sufficient to replenish the pool of inorganic carbon in the system that is converted to bicarbonate without aiming for excessively high pH levels. If the *CO*_2_ production is not sufficient for this purpose opting for 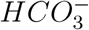 will be a better alternative. Inversely, if the *CO*_2_ concentration is high, using 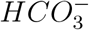 to control pH and alkalinity will be counterproductive as it will add to the reservoir of inorganic carbon that can be converted to *CO*_2_. Using a dosage system that accounts for the *CO*_2_ concentration in the system and combines both supplements thus presents a better alternative as shown in scenario 3.

Simulation results highlight the complexity of alkalinity control, illustrating how altering one parameter can change the entire system’s balance. Such changes can be non-intuitive, even to experts in the field. In their study, Summerfelt et al., (2015) [33] reported no significant differences in *CO*_2_ removal efficiency across various alkalinity treatments, contrary to their initial hypothesis. This experiment exclusively used *NaHCO*_3_ to regulate different alkalinity levels [33]. In contrast, Jafari et al., (2024) [19] found a significantly higher *CO*_2_ removal rate between nominal alkalinity 200 and 70. When simulating the experimental design from [19], the *CO*_2_ removal efficiency difference between nominal alkalinity 200 and 70 (Fig. 4) was more pronounced compared to simulation case 1, which only dosed 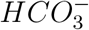 (Fig. 8). The authors in [33] also found no significant differences in *CO*_2_ concentra-tion between treatments, concurrent with the findings in simulation case 2, where *CO*_2_ concentration appeared to decrease mainly relative to fish growth rather than the alkalinity level when dosing with 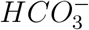 only (Fig. 7a). This highlights the model’s utility in exploring non-intuitive behaviours observed within the system.

While the model primarily focuses on alkalinity control, it may have broader applications. The dynRAS model’s ability to replicate feeding-induced patterns makes it a promising tool for designing water quality control routines. The model depicts large daily amplitude variations in most water quality parameters under a 12-hour intermittent feeding regime. This indicates that a parameter may be within the optimal range at sampling but exceed recommended levels hours later. Using such a model to approximate when and to what extent the concentrations of the metabolites will fluctuate over the day can guide RAS operators in tailoring their daily water quality control strategies.

Overall, the results suggest that standardizing alkalinity control to a single value and supplement may not be the best approach due to the dynamic nature of the system. Instead, adopting an adaptive control strategy is likely to be more effective. The complexity of the system dynamics in RAS makes establishing adaptive controls demanding without appropriate tools to understand and predict changes in the system dynamic. With future refinements, the dynRAS model presented here could act as a potent resource for the development of tools to work toward an adaptive control system for alkalinity and pH.

### 4.3. Toward smart RAS farming: tailoring the model to the system

The dynRAS model was implemented based on the structure and environmental setting of a pilot-scale RAS [19]. No parameter fitting was conducted as the aim was to reproduce observed patterns and inter-dependencies that can be generalized to RAS. As the model was implemented using an object-oriented approach, it can easily be modified and adapted to the topology of a specific system. However, although the rate laws used in the model to describe the system dynamic can be applied to other RAS, model parameters should be refined to the studied system to be used as a support for decision-making. When looking at the carbonate system in particular, the biological components play a major role in shaping these dynamics as illustrated by the model.

Tailoring models to specific systems is becoming more accessible with the rise of real-time monitoring systems that can continuously record parameters such as *CO*_2_, pH, temperature and *H*_2_*S* [20]. By combining these monitoring systems with machine learning, the model parameters could be continuously tuned to a particular system to optimize model predictions and refine water quality control.

Beyond parameter tuning, combining both data-driven and mechanistic models into hybrid models has already proven to enhance model accuracy for wastewater treatment plants [34]. While bottlenecks of mechanistic models can arise from the complexity of the modelled system, data-driven models suffer from a lack of interpretability and may struggle to accurately predict scenarios that they have not previously encountered. Using data-driven models in bridging the knowledge gaps of mechanistic models to enhance model predictions in new scenarios would mitigate the disadvantages of both approaches as suggested in [34]. Implementing this approach to RAS using the dynRAS model could provide a potent strategy to advance toward an autonomous control of alkalinity in RAS.

## 5. Conclusion

Through model support, we demonstrated the importance of understanding the system for optimal alkalinity control. RAS are inherently dynamic, and although environmental parameters are more strictly regulated with minimal external influences, we show that the system’s behaviour can vary considerably on day and week timescales, highlighting the importance of carefully designing water quality routines. As the biomass increases and *CO*_2_ production varies throughout the production cycle, adjusting the alkalinity level and selecting a base or buffer supplement based on the current system status, instead of committing to one option, proved to be more effective for system control. In light of this, using predictive models such as the dynRAS model to optimize alkalinity control might significantly enhance system efficiency. Although the model successfully reproduced the experimental data, it can be further refined. Combining machine learning techniques with mechanistic models like the dynRAS model presents an opportunity to enhance decision-making for water parameters management. This approach would not only optimizes environmental conditions for the fish but would also improve the overall system sustainability by reducing the need for excessive chemical interventions.

## CRediT authorship contribution statement

**Marie Aline Montjouridès:** Conceptualization, Formal analysis, Software, Investigation, Methodology, Writing - Original Draft, Writing - Review & Editing. **Susanna Röblitz:** Conceptualization, Methodology, Software, Investigation, Writing - Review & Editing, Supervision. **Håkon Dahle:** Conceptualization, Methodology, Investigation, Writing - Review & Editing, Supervision, Funding Acquisition

## Acknowledgments

The authors extend their gratitude to Anita Rønneset and the Fish Immunology Group for their help and valuable discussions in conceptualizing the experimental trial. Additional thanks go to Irene Roalkvam, Mark Powell, Melanie Andrews, and the RasLab staff for their technical assistance and support in designing the experiment. The authors also wish to thank Hanne Nilsen and the Norwegian Veterinary Institute in Bergen for providing fish measurement data. Additional thanks are due to Rune Sandvik for offering valuable insights into the industry’s needs during the project’s conceptualization. The authors are also grateful to SEARAS AS for providing the AquaSENSE™ system. Special thanks to Leila Jafari for the insightful discussions that helped improve the experiment, and to Olina Skorá for her assistance during the long sampling days. Finally, the authors thank Emily Olesin Denny for proofreading the paper.

## Data Availability

The data and the dynRAS model are available online at:

https://github.com/Marizauto/dynRAS.git

All the simulation scenario results can be reproduced using the provided model.

## Appendix A. Additional figure

**Fig. A.13.**
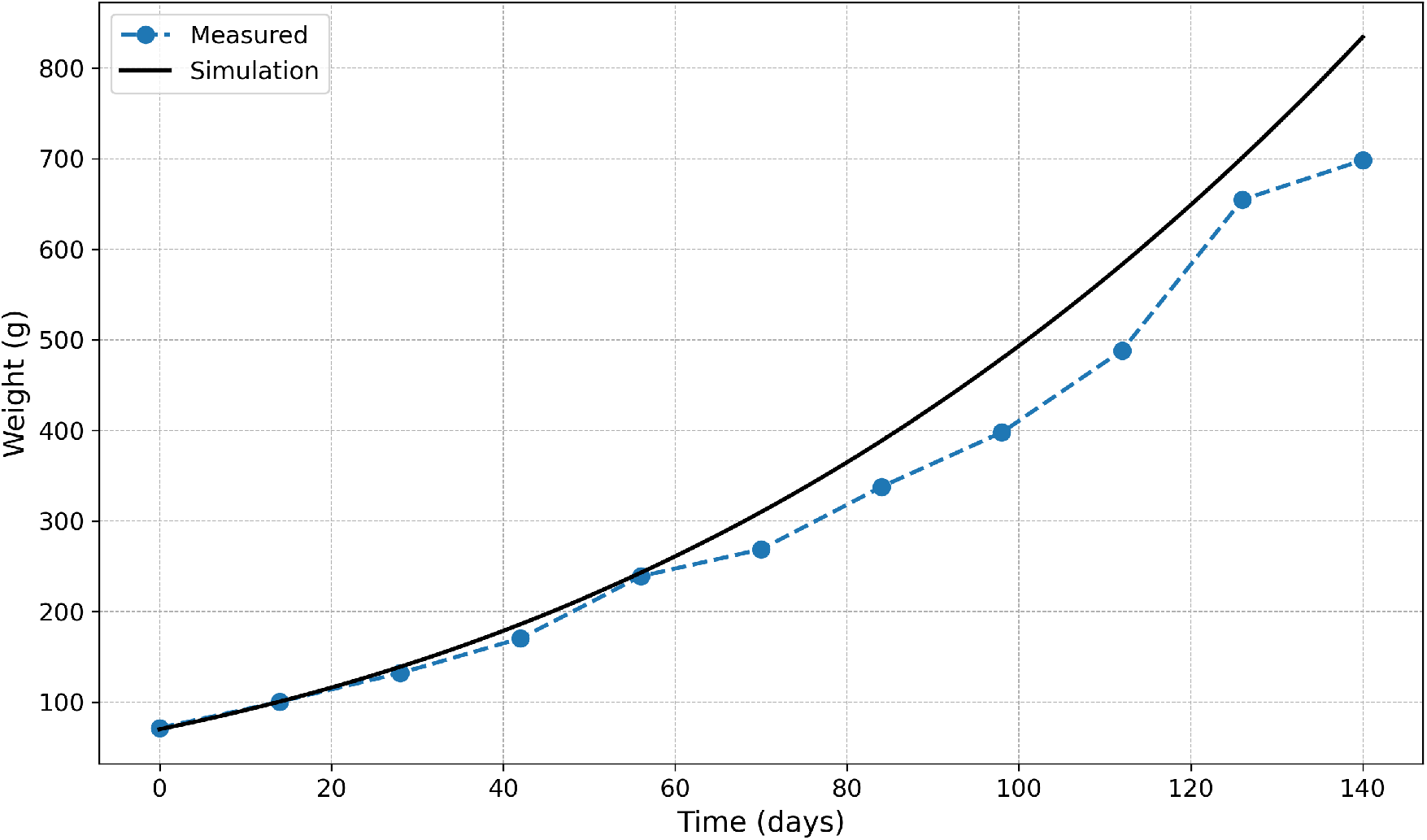
Mean fish weight for simulation results and measured data. The fish growth was the same for all simulation scenarios presented in the paper.

**Fig. A.14.**
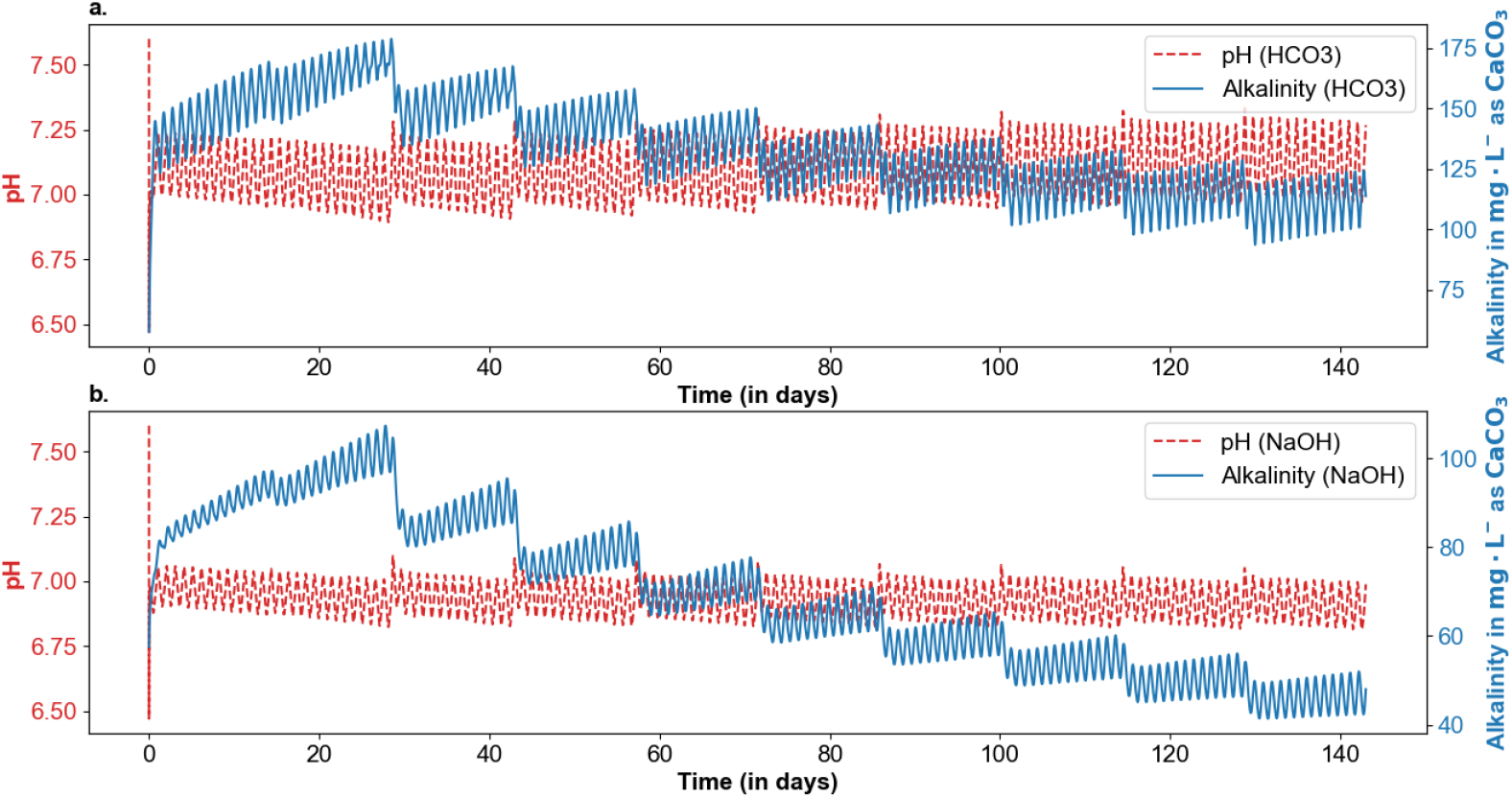
Model simulation for pH control using either a) 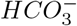 or b) NaOH. Blue lines represent the alkalinity in *mg* · *L*^−1^ as *CaCO*_3_ over time and red dashed lines the pH. The targeted threshold was 7 for both simulations. The simulations were run over 140 days with fish removed every 14 days to reduce the biomass to 50*kg* · *m*^−3^. As the fish grow, the *CO*_2_ production is slowly reduced and the alkalinity decreases.

## Appendix B. Full Equations

### Appendix B.1. Ordinary differential equations for the biofilter

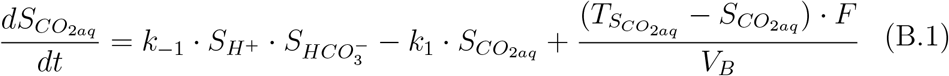

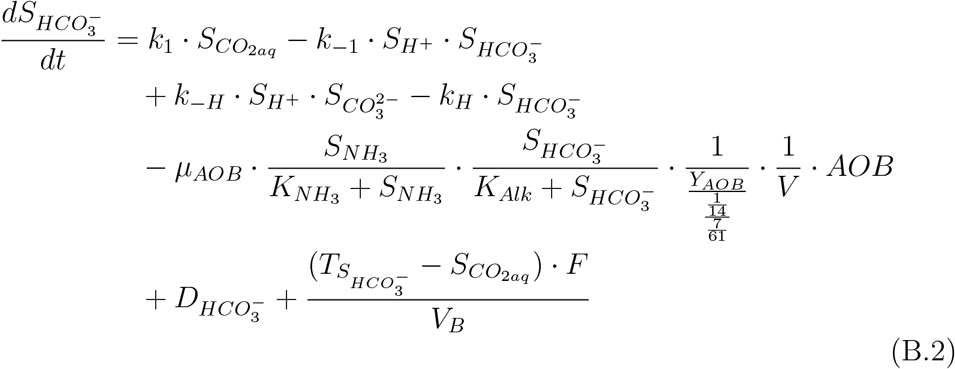

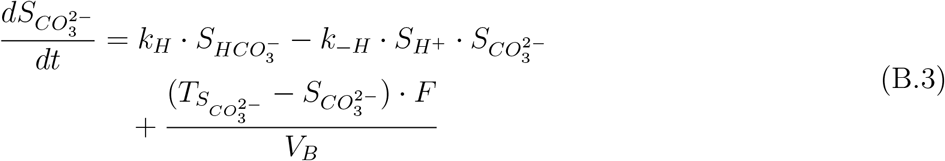

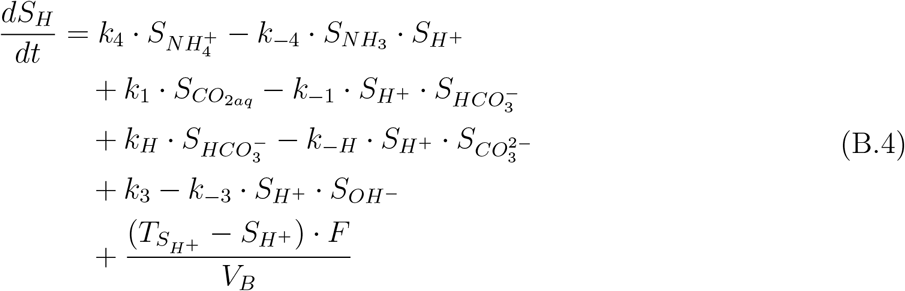

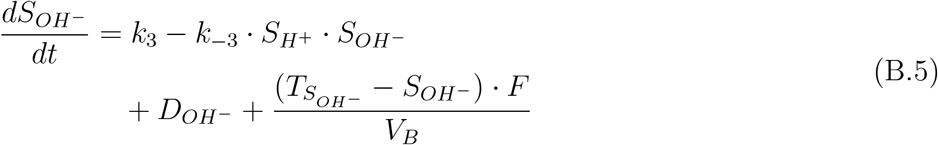

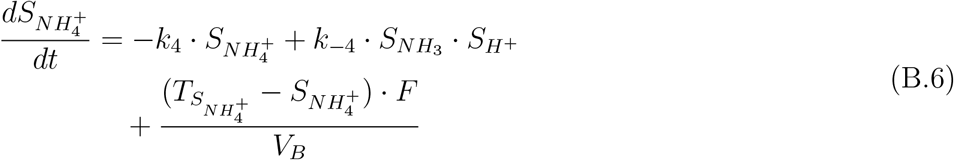

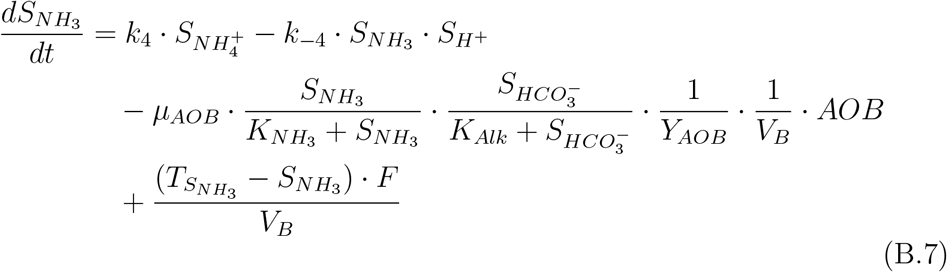

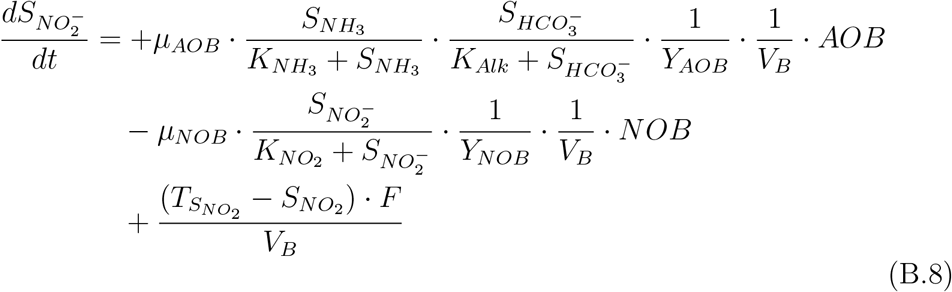

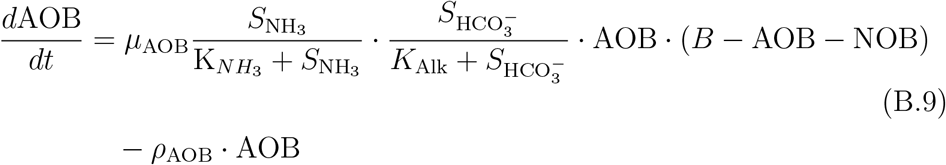

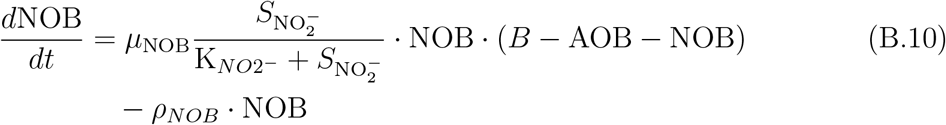

Here, 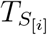 represent the concentration of the Species *i* coming from the Fish Tank, *V*_*B*_ is the volume of the biofilter, and 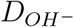 and 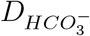 are the addition of *OH*^−^ and 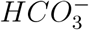 from the dosage system (eqs. (20),(21)).

### Appendix B.2. Ordinary differential equations for the Fish Tank

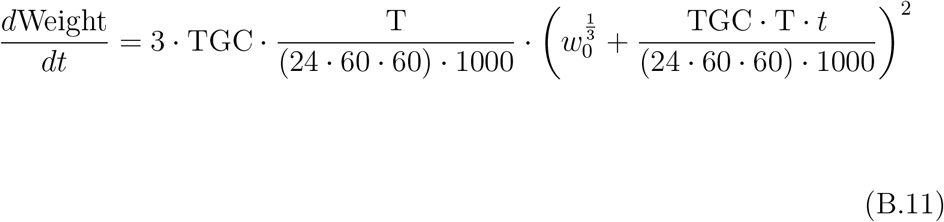

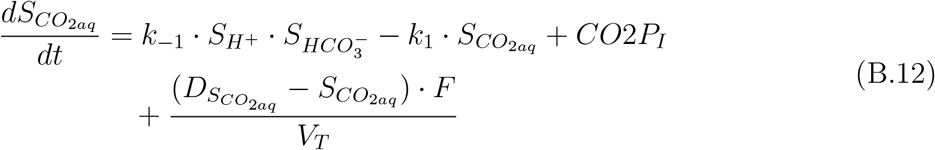

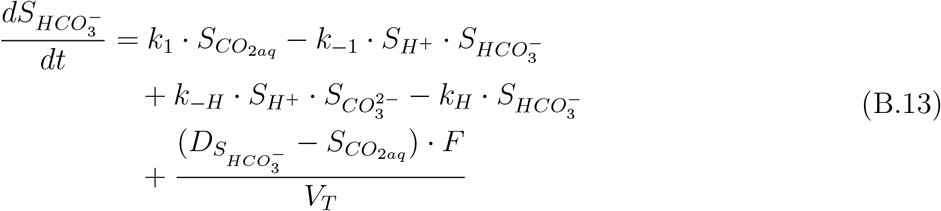

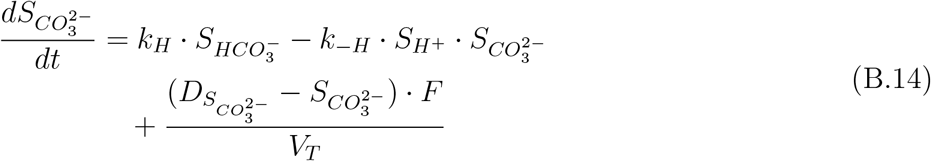

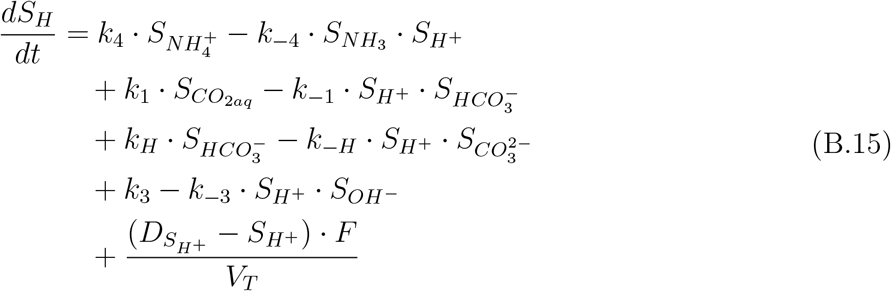

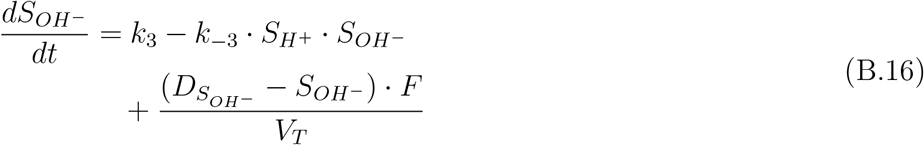

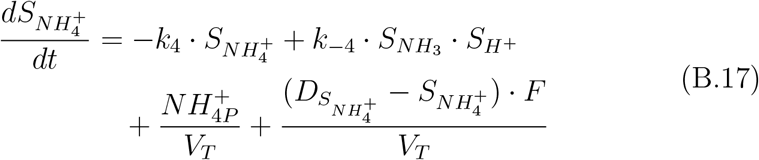

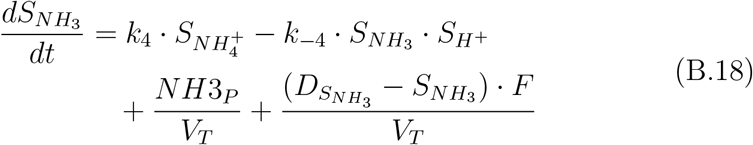

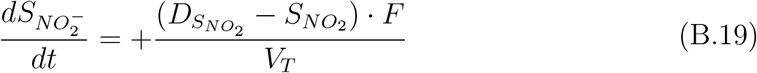

Here, *CO*2*P*_*I*_ is the production of *CO*_2_ from the fish (eq. (26)), *NH*3_*P*_ and 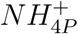 are the fraction of TAN excretion from feeding in the form of ammonium or ammonia in *mmol* · *s*^−1^ (eqs. (27), (28)), 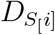 is the concentration of species *i* coming from the degasser, and *V*_*T*_ is the volume of the Tank.

### Appendix B.3. Ordinary differential equations for the Degasser

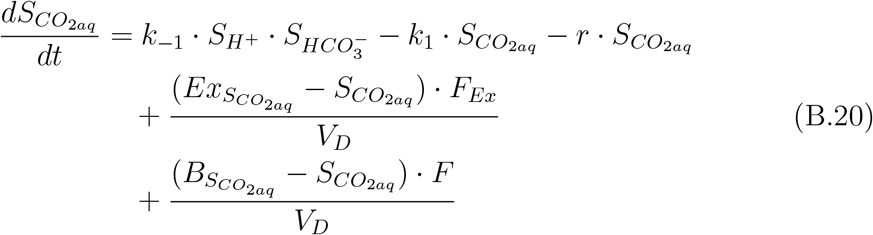

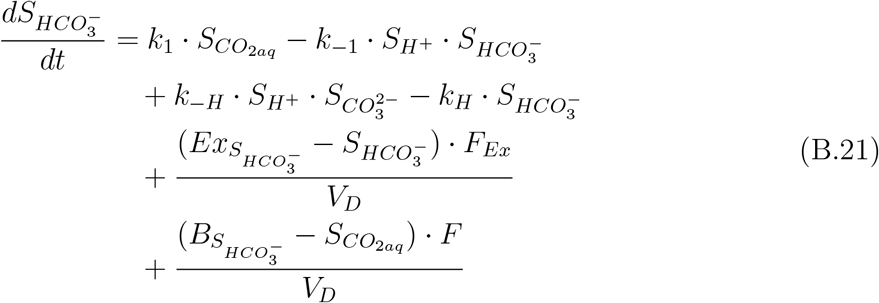

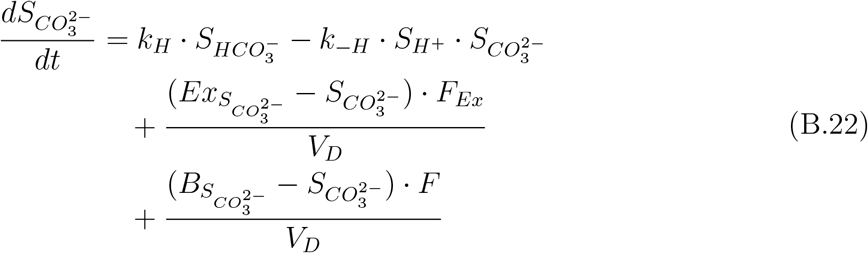

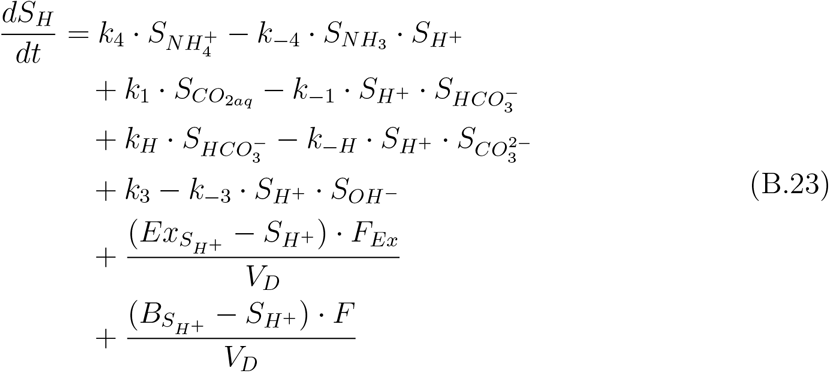

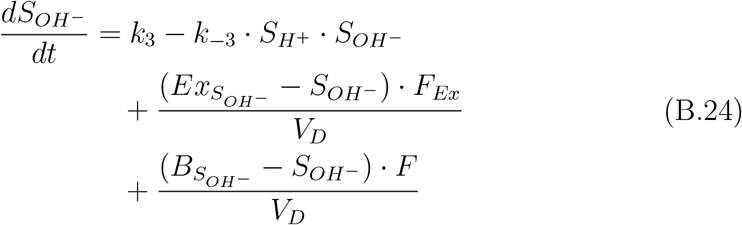

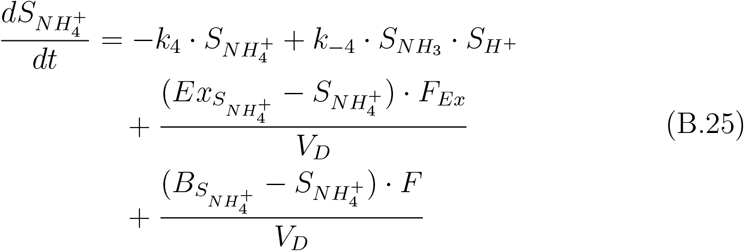

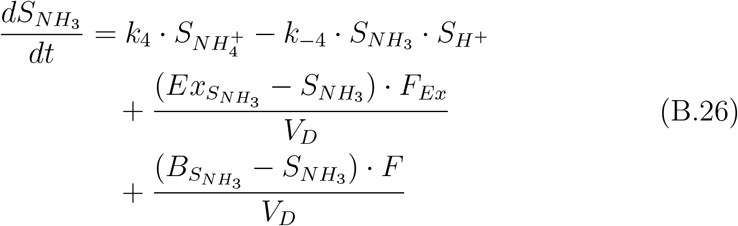

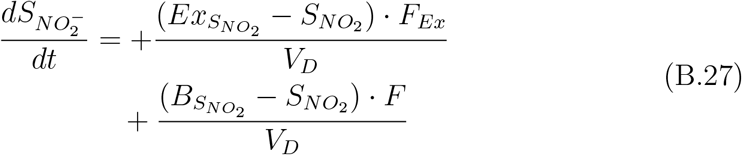

Here, 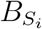 and 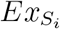 are the concentration of the species i in the biofilter and from the external source, respectively. *F*_*Ex*_ is the exchange rate in *L* · *s*^−1^, and *V*_*D*_ is the volume of the biofilter.

## References

[1] P. B. Bueno, D. Soto, Adaptation strategies of the aquaculture sector to the impacts of climate change, FAO (2017).

[2] R. Zhang, T. Chen, Y. Wang, M. Short, Systems approaches for sustainable fisheries: A comprehensive review and future perspectives, Sustainable production and consumption (2023).

[3] R. Xiao, Y. Wei, D. An, D. Li, X. Ta, Y. Wu, Q. Ren, A review on the research status and development trend of equipment in water treatment processes of recirculating aquaculture systems, Reviews in Aquaculture 11 (2019) 863–895.

[4] M. Badiola, D. Mendiola, J. Bostock, Recirculating aquaculture systems (ras) analysis: Main issues on management and future challenges, Aquacultural Engineering 51 (2012) 26–35.

[5] T. E. Wik, B. T. Lindén, P. I. Wramner, Integrated dynamic aquaculture and wastewater treatment modelling for recirculating aquaculture systems, Aquaculture 287 (2009) 361–370.

[6] E. Jiménez, J. Giménez, M. Ruano, J. Ferrer, J. Serralta, Effect of ph and nitrite concentration on nitrite oxidation rate, Bioresource technology 102 (2011) 8741–8747.

[7] S. Biesterfeld, G. Farmer, P. Russell, L. Figueroa, Effect of alkalinity type and concentration on nitrifying biofilm activity, Water environment research 75 (2003) 196–204.

[8] H. Pirouz Hamidi, The effect of pH and alkalinity on drinking water biofiltration performance, Ph.D. thesis, Carleton University, 2019.

[9] C. E. Boyd, C. S. Tucker, B. Somridhivej, Alkalinity and hardness: critical but elusive concepts in aquaculture, Journal of the World Aquaculture Society 47 (2016) 6–41.

[10] P. Lindholm-Lehto, Water quality monitoring in recirculating aquaculture systems, Aquaculture, Fish and Fisheries 3 (2023) 113–131.

[11] M. B. Timmons, J. M. Ebeling, F. W. Wheaton, S. T. Summerfelt, B. J. Vinci, Recirculating aquaculture systems, Cayuga Aqua Ventures, 2002.

[12] M. Henze, C. L. Grady, W. Gujer, G. Marais, T. Matsuo, A general model for single-sludge wastewater treatment systems., Water Research 21 (1987) 505–515.

[13] M. Henze, W. Gujer, T. Mino, M. Van Loosedrecht, Activated sludge models ASM1, ASM2, ASM2d and ASM3, IWA publishing, 2000.

[14] S. Pedersen, Simulation and Optimization of Recirculating Aquaculture Systems, Ph.D. thesis, Chalmers Tekniska Hogskola (Sweden), 2018.

[15] S. Pedersen, T. Wik, A comparison of topologies in recirculating aquaculture systems using simulation and optimization, Aquacultural engineering 89 (2020) 102059.

[16] A. M. dos Santos, L. F. Bernardino, K. J. Attramadal, S. Skogestad, Steady-state and dynamic model for recirculating aquaculture systems with ph included, Aquacultural Engineering 102 (2023) 102346.

[17] A. Bergheim, S. Fivelstad, Atlantic salmon (salmo salar l.) in aquaculture: Metabolic rate and water flow requirements, Salmon: Biology, Ecological Impacts and Economical Importance, 1st ed.; Woo, PTK, Noakes, DJ, Eds (2014) 155–173.

[18] I. Suzuki, U. Dular, S. Kwok, Ammonia or ammonium ion as substrate for oxidation by nitrosomonas europaea cells and extracts, Journal of Bacteriology 120 (1974) 556–558.

[19] L. Jafari, M. A. Montjouridès, C. D. Hosfeld, K. Attramadal, S. Fivelstad, H. Dahle, Biofilter and degasser performance at different alkalinity levels in a brackish water pilot scale recirculating aquaculture system (ras) for post-smolt atlantic salmon, Aquacultural Engineering 106 (2024) 102407.

[20] E. Lien, G. Valsvik, J. V. Nordstrand, V. Martinez, V. Rogne, O. Hafsås, S. Queralt, B. S. Fathi, M. Aga, The searas aquasense™ system: Realtime monitoring of h2s at sub μg/l levels in recirculating aquaculture systems (ras), Frontiers in Marine Science 9 (2022) 894414.

[21] K. G. Schulz, U. Riebesell, B. Rost, S. Thoms, R. Zeebe, Determination of the rate constants for the carbon dioxide to bicarbonate inter-conversion in ph-buffered seawater systems, Marine chemistry 100 (2006) 53–65.

[22] W. Stumm, J. J. Morgan, Aquatic chemistry: chemical equilibria and rates in natural waters, John Wiley & Sons, 2012.

[23] J. N. Weiss, The hill equation revisited: uses and misuses, The FASEB Journal 11 (1997) 835–841.

[24] K. S. Johnson, Carbon dioxide hydration and dehydration kinetics in seawater 1, Limnology and Oceanography 27 (1982) 849–855.

[25] H. Thorarensen, A. P. Farrell, The biological requirements for postsmolt atlantic salmon in closed-containment systems, Aquaculture 312 (2011) 1–14.

[26] A. Berg, A. Danielsberg, A. Seland, T. Sigholt, Oxygen demand for postsmolt atlantic salmon (salmo salar l.), in: Fish farming technology, CRC Press, 1993, pp. 297–300.

[27] J. A. Grøttum, T. Sigholt, A model for oxygen consumption of atlantic salmon (salmo salar) based on measurements of individual fish in a tunnel respirometer, Aquacultural engineering 17 (1998) 241–251.

[28] E. Eliason, A. Farrell, Oxygen uptake in pacific salmon oncorhynchus spp.: when ecology and physiology meet, Journal of Fish Biology 88 (2016) 359–388.

[29] A. Bergheim, E. A. Seymour, S. Sanni, T. Tyvold, S. Fivelstad, Measurements of oxygen consumption and ammonia excretion of atlantic salmon (salmo salar l.) in commercial-scale, single-pass freshwater and seawater landbased culture systems, Aquacultural Engineering 10 (1991) 251–267.

[30] R. G. Bates, G. D. Pinching, Acidic dissociation constant of ammonium ion at 0 to 50 c, and the base strength of ammonia, J. Res. Natl. Bur. Stand 42 (1949) 419–430.

[31] M. Eigen, Proton transfer, acid-base catalysis, and enzymatic hydrolysis. part i: elementary processes, Angewandte Chemie International Edition in English 3 (1964) 1–19.

[32] A. V. Bandura, S. N. Lvov, The ionization constant of water over wide ranges of temperature and density, Journal of physical and chemical reference data 35 (2006) 15–30.

[33] S. T. Summerfelt, A. Zühlke, J. Kolarevic, B. K. M. Reiten, R. Selset, X. Gutierrez, B. F. Terjesen, Effects of alkalinity on ammonia removal, carbon dioxide stripping, and system ph in semi-commercial scale water recirculating aquaculture systems operated with moving bed bioreactors, Aquacultural Engineering 65 (2015) 46–54.

[34] W. Quaghebeur, E. Torfs, B. De Baets, I. Nopens, Hybrid differential equations: integrating mechanistic and data-driven techniques for modelling of water systems, Water Research 213 (2022) 118166.

